# Endometriosis patient-derived small extracellular vesicles carry unique immune, proteomic and lipidomic signatures associated with mild and severe endometriosis

**DOI:** 10.64898/2026.08.13.744471

**Authors:** Jaelis P. Holmes, Katherine B. Zutautas, Danielle J. Sisnett, Donya Hayati, Olga Bougie, Bruce A. Lessey, Chandrakant Tayade

**Author notes:** **Corresponding author:** Dr. Chandrakant Tayade1, DVM, PhD, Department of Biomedical and Molecular Sciences, Queen’s University, Kingston, ON, Canada, K7L 3N6.

## Abstract

Endometriosis (EM) is a heterogeneous, gynecological inflammatory disease affecting over 200 million individuals worldwide, yet the mechanisms underlying lesion establishment, progression, and recurrence remain incompletely understood. Small extracellular vesicles (sEVs) mediate intercellular communication through the transfer of proteins, lipids, and nucleic acids reflective of their cellular origin; however, stage- and tissue-specific sEV signatures remain poorly defined. Here, we characterized the molecular and functional landscape of EM-derived sEVs across disease stages and biological sources. sEVs isolated from eutopic endometrium, ectopic lesions, peritoneal fluid, and plasma from mild- and severe-stage EM patients and healthy controls were analyzed by surface marker profiling, proteomics, lipidomics, and integrated multi-omics, with functional effects assessed in human uterine microvascular endothelial cells. sEV composition varied by disease stage and sample type, with EM lesion-derived sEVs demonstrating stage-dependent loss of epithelial-associated markers and enrichment of immune-associated signatures, while EM plasma-derived sEVs exhibited altered adhesion- and platelet-associated profiles. Integrated multi-omics identified coordinated programs associated with immune adaptation, extracellular matrix organization, epithelial remodeling, vascular signaling, oxidative stress, and metabolic adaptation. Functionally, sEVs derived from severe endometriotic lesions exhibited enhanced uptake and mitochondrial localization in endothelial cells and promoted angiogenic activity. Our findings establish sEVs as dynamic mediators of EM disease progression and demonstrate that integrated sEV profiling provides a framework for understanding EM heterogeneity and identifying candidate biomarkers and therapeutic targets.

## 1. Introduction

Endometriosis (EM) is a chronic, estrogen-dependent inflammatory disease characterized by the presence of endometrium-like tissue outside the uterine cavity, primarily on the pelvic peritoneum and ovaries.^1,2^ Affecting approximately 6–10% of reproductive-age women worldwide, EM is associated with chronic pelvic pain, dysmenorrhea, dyspareunia, and infertility, substantially reducing quality of life and imposing a significant socioeconomic burden.^1,2^ Disease severity is clinically classified into four stages (I–IV) according to the revised American Society for Reproductive Medicine (rASRM) staging system, with stages I– II generally considered mild disease and stages III–IV considered severe disease.^1^ However, symptom presentation and disease progression remain highly heterogeneous between patients, reflecting the complex and incompletely understood pathogenesis of EM. Increasing evidence implicates immune dysregulation, chronic inflammation, and altered cellular communication in lesion establishment and disease progression.^3,4^ Notably, mild and severe disease stages are thought to reflect distinct pathological states, with mild disease associated with adhesion and survival within the peritoneal environment, and severe disease characterized by persistent inflammation, angiogenesis, and progressive tissue remodeling and fibrosis.^3,4^ Despite advances in diagnosis and clinical management, significant diagnostic delays and limited non-invasive biomarkers continue to challenge effective disease detection and characterization, highlighting the need for improved understanding of EM-associated molecular and cellular mechanisms.^2,5^

Extracellular vesicles (EVs) have emerged as important mediators of intercellular communication in both physiological and pathological conditions.^6,7^ EVs are lipid membrane-bound particles released by nearly all cell types and carry diverse bioactive cargo, including proteins, lipids, and nucleic acids, reflective of their cell of origin.^6,7^ They differ in size, biogenesis, and biological function and are classified into distinct subtypes, including apoptotic bodies, microvesicles, and small extracellular vesicles (sEVs). Among these, sEVs are abundant in the uterine microenvironment and are increasingly recognized for their roles in immune modulation, angiogenesis, proliferation, migration, and inflammatory signaling.^6–11^ These small vesicles, typically ranging from 50–250 nm, encompass populations commonly associated with exosomes and small microvesicles.^6,7^ Through receptor-ligand interactions, membrane transfer, and delivery of bioactive cargo, sEVs can alter cellular phenotype and function both locally and systemically.^7^ Importantly, sEVs are emerging as carriers of disease-associated molecular signatures, making them attractive candidates for investigating mechanisms of disease progression and identifying minimally invasive biomarkers.

In the context of EM, sEVs exhibit disease-associated alterations to their molecular cargo, including differential expression of miRNAs, lncRNAs, proteins, and lipids relative to healthy controls.^12–14^ Among the most extensively studied sEV cargo are miRNAs, including members of the let-7 family, miR-30d-5p, miR-23a, miR-143, and miR-320a, which have been implicated in inflammatory, proliferative, and angiogenic signaling pathways in EM.^13–17^ Similar miRNA alterations have also been detected in the plasma of EM patients.^13,15^ Complimenting these observations, recent work using bead-based flow cytometric profiling of sEV surface epitopes in a patient-derived endometrial epithelial organoid model demonstrated stage-dependent remodeling of sEV phenotypes, with severe-stage EM-derived sEVs exhibiting increased immune-associated surface markers and reduced stem cell-associated markers relative to control and mild-stage disease.^18^ Together, these findings support a role for sEVs in EM pathophysiology. However, most studies have focused on individual sEV cargo classes or isolated biofluids, leaving the extent to which sEV composition varies across EM disease stages and matched tissue compartments poorly understood. Moreover, whether these molecular signatures reflect coordinated alterations across the local lesion microenvironment and systemic circulation remains unclear.

Building on this gap, we aimed to define stage- and tissue-specific sEV signatures in EM through integrated surface marker, proteomic, and importantly lipidomic profiling of sEVs isolated from eutopic endometrium, ectopic lesions, plasma, and peritoneal fluid. Integrating lipidomic with proteomic profiling enabled characterization of coordinated molecular remodeling across complementary sEV cargo classes, providing insight into immune, metabolic, extracellular matrix (ECM), and vascular remodeling signatures associated with EM progression. Surface marker profiling revealed stage-dependent shifts in immune-, adhesive-, and stemness-associated sEV phenotypes, while proteomic and lipidomic analyses uncovered distinct molecular programs linked to lesion establishment and severe disease remodeling. As angiogenesis and vascular remodeling are essential features of EM lesion establishment and persistence, and were among the dominant pathways identified through our integrated analyses, we further evaluated whether stage-specific lesion-derived sEVs differentially interact with endometrial endothelial cells by assessing sEV uptake, cytokine secretion, and angiogenic capacity. Our findings provide insight into stage- and tissue-dependent molecular signatures and functional activity of sEVs in EM, supporting their potential role(s) in disease progression and their utility as minimally invasive indicators of EM-associated pathophysiology.

## 2. Methods

### 2.1 Study approval and ethics

The Queen’s University Health Sciences and Affiliated Teaching Hospitals Research Ethics Board approved all methodologies used in this study (OBGY-229-11 and ANAT-029-09). All participants recruited from gynecology clinics at Kingston Health Sciences Centre (KHSC) provided written informed consent prior to sample collection. Samples used for lipidomic analyses were obtained from an independent cohort collected through Greenville Hospital System (Greenville, South Carolina, USA) and Kingston General Hospital (KGH), following institutional research ethics approval and written informed consent from all participants.

The primary KHSC cohort was used for extracellular vesicle characterization and functional analyses consisting of matched samples from mild-stage (n=5) and severe-stage EM patients (n=7), including ectopic lesions, eutopic endometrium, peritoneal fluid (PF), and plasma, as well as healthy control plasma samples (n=8).^19^ An independent cohort was utilized for lipidomic profiling (**Table 1**), consisting of EM patient plasma samples (mild, n=8; severe, n=6), eutopic endometrium samples (n=4), and ectopic lesion samples (mild, n=4; severe, n=4) collected from Greenville Hospital System, with healthy control plasma samples (n=6) obtained through KGH. Across both cohorts, participants met eligibility criteria of being 28–50 years of age, having a uterus and at least one ovary, and undergoing scheduled excision surgery within the study period. Individuals with EM received a confirmed diagnosis of EM by histological evidence, suspicious clinical history, and/or radiographic evidence consistent with EM, prior to surgery. Inclusion and exclusion criteria, sample processing procedures, and storage conditions were consistent between cohorts.

**Table 1.**
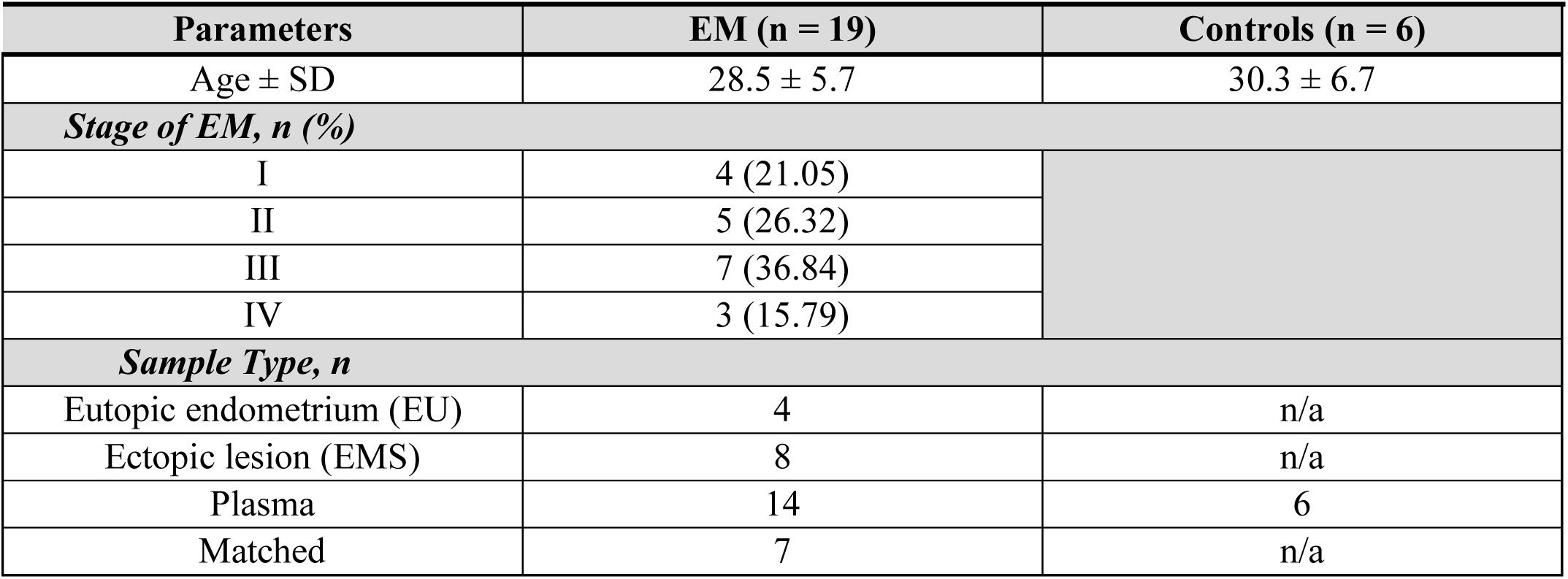
Clinical characteristics of independent cohort used for lipidomic analyses.

### 2.2 Sample collection from EM patients and control women

For proteomic profiling and sEV characterization cohort, matched human eutopic endometrium and endometriotic lesion biopsies, and PF samples were collected from patients undergoing laparoscopic EM excision surgery. Histopathological analysis of excised lesions was performed to confirm EM diagnosis.

Peripheral blood was collected prior to surgery. For the independent lipidomic cohort, eutopic endometrial samples were obtained by Pipelle biopsy during surgery, while ectopic lesions and PF were collected during laparoscopic excision procedures. All samples were processed and stored according to standardized procedures, with tissue samples snap-frozen in liquid nitrogen and stored at −80°C until further analysis. PF samples were similarly stored at −80°C, and plasma was isolated from peripheral blood as described below. EM stage was determined intraoperatively by the attending surgeon according to the revised American Society for Reproductive Medicine (rASRM) classification criteria.^1^ Clinical and demographic characteristics of matched EM patient cohort used for sEV characterization, proteomic profiling, and functional analyses, including age, pathology findings, co-existing pathologies, and disease stage (I–IV), were previously reported by our group. ^19^ Control participants (plasma only) had no clinical indicators of EM or other gynecological conditions, including infertility, pelvic inflammatory disease, or chronic pelvic pain. Clinical characteristics of independent cohort used for lipidomic analyses followed similar inclusion criteria (**Table 1**).

### 2.3 Plasma isolation and PF collection

Peripheral blood samples from control and EM patients were diluted 1:1 with 2% FACS buffer (PBS supplemented with 2% FBS) prior to processing using Lymphoprep density gradient medium (18060; STEMCELL Technologies) and SepMate™-50 tubes (85450; STEMCELL Technologies), according to the manufacturer’s instructions. Following density gradient centrifugation, the plasma fraction was carefully aspirated and immediately stored at −80°C until further use. PF was processed as per our previous publication.^19^

### 2.4 Tissue protein extract and quantification

Approximately 50 mg of frozen eutopic endometrium (EU)- or endometriotic lesion (EMS)-derived tissue was obtained from each sample while maintaining samples on dry ice to prevent thawing. Tissue samples were transferred into PowerBead tubes (13112-50; Qiagen) preloaded with T-PER™ Tissue Protein Extraction Reagent (78510; Thermo Fisher Scientific) supplemented with protease inhibitor cocktail (535140-1ML; Sigma-Aldrich) at a 1:100 ratio. Samples were then homogenized using an Omni Bead Ruptor 24 (Omni International) for two cycles of 20 s at 5.5 m/s with a 20 s interval between cycles. Homogenization parameters were adjusted as required based on tissue size and fibrous composition. Following homogenization, samples were centrifuged at 10,000 × g for 5 min at 4°C to remove insoluble debris. The resulting protein-containing supernatant was collected on ice prior to protein quantification and storage at −80°C until further analysis. Briefly, protein concentration was determined using a bicinchoninic acid (BCA) assay (23227; Thermo Fisher Scientific), as per manufacturer’s guidelines.

### 2.5 Isolation of EVs, including sEVs, from patient plasma, tissues, and PF

sEVs were isolated from plasma, PF, and tissue samples obtained from EM patients and healthy controls using size exclusion chromatography (SEC). IZON qEV columns (IC10-35 for plasma and PF samples; ICO-35 for tissue samples; IZON Science, Christchurch, New Zealand) were used according to the manufacturer’s instructions. Fractions 7–9, corresponding to sEV-enriched fractions, were collected for downstream analyses. All EV isolation procedures were performed in accordance with the MISEV2025 guidelines.^20^ Samples underwent additional filtration through a 0.22-μm filter to remove potential contaminants. The quality of sEVs isolated using SEC was compared with ultracentrifugation-based isolation methods and was determined to be superior with respect to purity and yield. Isolated sEV pellets were resuspended in 150 μL of PBS (10010023, Thermo Fisher Scientific) and either used immediately for downstream applications or stored at –80°C until further use. sEV samples were subjected to no more than 2 freeze-thaw cycles. The isolated sEV preparations described above, derived from matched patient eutopic endometrium, ectopic lesions, plasma, and PF samples from individuals with mild (n=5) and severe (n=7) EM, as well as plasma-derived sEVs from healthy controls (n=8), were used for downstream characterization, MACSPlex surface marker analysis, and proteomic profiling.

### 2.6 Transmission electron microscopy analysis of sEVs

Isolated sEVs were characterized by transmission electron microscopy (TEM). sEVs resuspended in 150μL PBS were fixed in 2.5% glutaraldehyde for 5 min and negatively stained with UranyLess (22409; Electron Microscopy Sciences) for 2 min. Samples were transferred onto 200-mesh Formvar-coated copper grids and incubated for 10 min. Grids were analyzed using a Talos F200i transmission electron microscope operated at 200 keV (Thermo Fisher Scientific) by trained electron microscopy specialists at the Queen’s University Cardiopulmonary Unit. Detected sEVs were subjected to morphometric analysis.

### 2.7 Nanoparticle tracking analysis of sEVs

Isolated sEVs were analyzed using a ZetaView nanoparticle tracking analyzer (Particle Metrix). Instrument alignment was performed using 100 nm reference standard polystyrene beads (3100A; Thermo Fisher Scientific). sEV samples were diluted 1:100 in PBS and analyzed at a sensitivity setting of 70–80 across 11 predefined camera positions. According to manufacturer guidelines, samples with fewer than 8/11 valid position reads or fewer than 500 tracked particles were excluded from analysis. Particles measuring <30 nm were considered background artifacts and excluded from quantification.

### 2.8 Detection and characterization of sEVs with MACsPlex analysis

Isolated sEVs were characterized using the MACSPlex Human EV Kit (130-108-813; Miltenyi Biotec). Positive marker expression was defined as fluorescence intensity exceeding the corresponding isotype control threshold. For each sample, 15 μg of EV-associated protein, quantified using a BCA protein assay (A65453; Thermo Fisher Scientific), was diluted in 120 μL MACSPlex buffer according to the manufacturer’s recommendations. Samples were acquired using a CytoFLEX S flow cytometer (Beckman Coulter) and analyzed using FlowJo™ software (BD Life Sciences, v10). This bead-based flow cytometry platform evaluates 37 EV surface markers alongside two isotype controls. Relative marker expression was calculated as normalized median fluorescence intensity (nMFI). Plasma-derived sEVs were normalized to the average signal of CD63 and CD81, whereas PF- and tissue-derived sEVs were normalized to the average signal of CD9, CD63, and CD81. The selection of normalization markers was based on consistent tetraspanin detection patterns within each sample matrix.

### 2.9 sEV protein isolation

Total protein was extracted from isolated sEVs, using a Total Exosome RNA & Protein Isolation Kit (4478545; Thermo Fisher Scientific) according to the manufacturer’s instructions. Protein concentration was quantified using a BCA protein assay (A65453; Thermo Fisher Scientific). Protein fractions were immediately stored at –80°C until downstream analyses.

### 2.10 Proteomic profiling of sEVs by mass spectrometry

Proteomic profiling was performed by the Proteomics Platform at the Research Institute of the McGill University Health Centre (RI-MUHC; Montreal, QC, Canada). Isolated sEV proteins were desalted by dialysis against 10 mM ammonium bicarbonate and subsequently digested with trypsin (20 μg) at 37°C for 18 h.

Resulting peptide mixtures were analyzed by liquid chromatography-tandem mass spectrometry (LC-MS/MS) using an Orbitrap Astral mass spectrometer (Thermo Fisher Scientific). Sample preparation, data acquisition, and initial processing were completed by the RI-MUHC Proteomics Platform according to standard facility methodologies. MS/MS spectra were acquired in positive ion mode and processed for downstream proteomic analysis. MS/MS-derived proteomic datasets were analyzed using DIA-NN and Spectronaut databases (v0.11) through the Analyst Suites platform.^21^ Protein abundance values were normalized using variance-stabilizing normalization (VSN), and missing values were not imputed. Differentially expressed proteins were identified using predefined statistical thresholds, including a false discovery rate (FDR) of 1%, adjusted p-value < 0.05, and a fold-change cutoff of |log2FC| ≥ 0.5. Functional enrichment and pathway analyses were performed using Integrated Pathway Analyst software, while obaDIA was used for protein functional annotation and Gene Ontology (GO) and Kyoto Encyclopedia of Genes and Genomes (KEGG) enrichment analyses.

### 2.11 Lipidomic profiling of sEVs by mass spectrometry

Lipidomic profiling was performed by Creative Proteomics (Shirley, New York, USA). sEVs isolated from matched patient eutopic endometrium (n=4), ectopic lesions from patients with mild (n = 4) and severe (n = 4) EM, alongside plasma from mild (n=8) and severe (n=6) patients, and healthy controls (n =6), were separated into 50 μL aliquots, followed by the addition of 1.5 mL chloroform:methanol (2:1, v/v) and 0.5 mL ultrapure water. Samples were vortexed and centrifuged to induce phase separation. The lower organic phase was carefully collected and dried under nitrogen gas. Dried lipid extracts were resuspended in isopropanol:methanol (1:1, v/v), and 5 μL lysophosphatidylcholine (LPC)(12:0) internal standard was added prior to analysis. Samples were subsequently centrifuged at 12,000 rpm for 10 min at 4°C and the resulting supernatant was collected for LC-MS analysis, which was performed using an ACQUITY UPLC system coupled to a Q Exactive mass spectrometer (Thermo Fisher Scientific). Chromatographic separation was achieved using an ACQUITY UPLC BEH C18 column (100 × 2.1 mm, 1.7 μm particle size; Waters). The mobile phase consisted of solvent A [60% acetonitrile (ACN), 40% H₂O, and 10 mM ammonium formate] and solvent B [10% ACN, 90% isopropanol, and 10 mM ammonium formate]. Gradient elution was performed as follows: 0–1 min, 30% B; 1–10.5 min, 30–100% B; 10.5–12.5 min, 100% B; 12.5–12.51 min, 100–30% B; and 12.51–16 min, 30% B. The mobile phase flow rate was maintained at 0.3 mL/min. Mass spectrometry data were acquired in both positive and negative electrospray ionization modes using optimized instrument parameters.

### 2.12 Integration of proteomic and lipidomic profiling of sEVs

Raw proteomic and lipidomic data files generated from the analyses described above were uploaded to the publicly available OmicsAnalyst platform for multi-omics integration.^22,23^ Datasets were formatted according to platform requirements and subjected to feature scaling and normalization within the OmicsAnalyst workflow. For tissue-derived sEV analyses, eutopic endometrium samples were designated as the reference group, whereas healthy control plasma samples served as the reference for plasma-derived sEV analyses. FDR correction was applied throughout the OmicsAnalyst workflow. Multiple Co-Inertia Analysis (MCIA) was performed to integrate proteomic and lipidomic datasets. Integrated features were further analyzed by Reactome pathway enrichment and causal discovery analyses to identify coordinated molecular relationships. Hierarchical clustering heatmaps were generated using Spectrum clustering, and integrated protein-lipid correlation networks were annotated using the STRING database.

### 2.13 Cell culture

For confocal microscopy-based sEV internalization and subcellular localization analyses, pooled sEVs isolated from ectopic lesions from individuals with mild (n=3) and severe (n=3) EM were used. For *in vitro* functional assays, sEVs isolated from matched patient ectopic lesions and eutopic endometrium from individuals with mild (n=5) and severe (n=7) EM were used as experimental sEV preparations. The same pooled sEV preparations were maintained across respective experimental assays to ensure consistency between experiments. Human uterine microvascular endothelial cells (HUtMEC; C-12295; PromoCell) were cultured according to manufacturer-provided protocols in Endothelial Cell Growth Medium MV (C-22020; PromoCell), a low-serum (5% v/v) medium supplemented with fetal calf serum (0.05 ml/ml), endothelial cell growth supplement (0.004 ml/ml), recombinant human epidermal growth factor (10 ng/ml), heparin (90 µg/ml), and hydrocortisone (1 µg/ml). Only passages 4-6 were used in experiments. Cellular morphology and proliferative characteristics were routinely monitored throughout passaging to ensure maintenance of phenotype and cellular integrity.

### 2.14 Point-scanning confocal microscopy analysis of sEV uptake and subcellular localization

sEVs were fluorescently labeled with 2 μM MemGlow™ 488 (MG01; Cytoskeleton Inc.) at 37°C for 10 min according to the manufacturer’s protocols Following staining, sEVs were washed using 100 kDa Amicon Ultra centrifugal filters (UFC5100; Sigma-Aldrich) to remove excess dye. HUtMEC cells were seeded at a density of 5 × 10⁴ cells/well in μ-Plate 24-well glass-bottom plates (82427; ibidi) and cultured for 24 h prior to treatment. For assessment of sEV uptake, cells were stained with 1 µg/mL Hoechst nuclear dye (62249; Thermo Fisher Scientific) and 1 μM CellTrace™ BODIPY® TR methyl ester cytoplasmic dye (C34556; Thermo Fisher Scientific) at 37°C for 30 min according to the manufacturer’s protocols. For assessment of sEV subcellular localization, cells were stained with 1 µg/mL Hoechst nuclear dye (62249; Thermo Fisher Scientific) and 1 μM MitoTracker™ Deep Red FM mitochondrial dye (M22426; Thermo Fisher Scientific) under the same conditions. Fluorescently labeled sEVs were resuspended in respective cell culture media and added to HUtMECs at a dose of 1 × 10⁴ sEVs/cell for 10 h.

Live-cell imaging was performed using a MICA point-scanning confocal microscope (Leica Microsystems) maintained at 5% CO₂, 60% humidity, and 37°C. Image acquisition was initiated immediately following sEV treatment and continued at 2 h intervals over the 10 h incubation period. Images were acquired at 60× magnification using a water-immersion objective lens and processed using LAS X Analyst Suite (Leica Microsystems) and ImageJ software (NIH).

### 2.15 Proliferation and apoptosis assays

HUtMEC cells were seeded in 96-well plates (163320; Thermo Fisher Scientific) and cultured for 24 h prior to sEV treatment. sEVs were resuspended in endothelial cell culture media and added to cells at a dose of 1 × 10⁴ sEVs/cell for 24 h. To assess cell proliferation and viability, cells were incubated with 10 μL WST-1 reagent (5015944001; Sigma-Millipore) for 2 h according to the manufacturer’s instructions. Cellular metabolic activity was quantified by measuring absorbance at 450 nm with a 650 nm reference wavelength using a SpectraMax iD3 plate reader (Molecular Devices). To evaluate apoptosis, cells were treated with 100 μL Caspase-Glo® 3/7 reagent (G8091; Promega) for 3 h protected from light, and caspase-3/7 activity was quantified by luminescence using the SpectraMax iD3 platform according to the manufacturer’s instructions.

All conditions were analyzed using 10–15 technical replicates.

### 2.16 Multiplex cytokine analysis

In parallel with *in vitro* functional assays, 100 μL of conditioned media was collected from HUtMEC cells 12 h following treatment with pooled sEVs. Cell-only controls were included. Inflammatory cytokine and chemokine profiling were performed using a commercially available Human Cytokine/Chemokine Panel A 48-Plex Discovery Assay® Array (Eve Technologies; HD48A) on the Luminex xMAP platform (Bio-Rad). All conditions were analyzed in three technical replicates.

### 2.17 Endothelial tube formation assay

Endothelial tube formation assays were performed using 15 well μ-Slide plates (81506; ibidi) according to the manufacturer’s instructions. Growth factor-reduced, phenol red-free Matrigel (356221; Corning) was dispensed into each well and allowed to polymerize at 37°C for 45 min. HUtMECs were harvested using 0.25% trypsin-EDTA, seeded onto the polymerized Matrigel at a density of 1 × 10⁴ cells/well, and treated with sEVs at a dose of 1 × 10⁴ sEVs/cell. Controls included PBS vehicle and recombinant human VEGF-A (25 ng/mL; MA116629; Thermo Fisher Scientific). Cells were incubated at 37°C and imaged at 12 h using a MICA widefield imaging system (Leica Microsystems). Two images were acquired per well from standardized regions along the well midline to ensure consistent image sampling across conditions. All conditions were analyzed in three technical replicates. Tube formation parameters, including number of tubes, total tube length, and branching points, were analyzed using the automated online platform WimTube (Wimasis GmbH, Munich, Germany).

### 2.18 Statistics

Statistical analyses were performed using GraphPad Prism software (v11). Data are presented as mean ± standard deviation (SD). Normality and homogeneity of variance were assessed prior to statistical testing. Outliers were identified using the ROUT method (Q = 1%) in GraphPad Prism; no outliers were removed unless indicated in respective figure captions. Statistical comparisons were performed using one-way analysis of variance (one-way ANOVA), two-way analysis of variance (two-way ANOVA), or repeated-measures one-way ANOVA, as appropriate based on experimental design, followed by Tukey’s multiple comparisons test for post hoc analysis. For experiments involving repeated measurements, repeated-measures analyses were applied. A p-value < 0.05 was considered statistically significant.

## 3. Results

### 3.1 EM-derived sEVs display tissue- and stage-specific biomolecular and surface phenotypic profiles

To elucidate stage- and tissue-specific alterations in sEVs in EM, we characterized sEV-enriched fractions from matched EM patient samples, and plasma samples from disease-free controls for comparison. sEVs were isolated by SEC across all sample types and disease stages (**Fig. 1a-n**). NTA revealed no significant differences in median particle size between biological compartments, with plasma-sEVs exhibiting consistent particle size distributions across systemic samples and PF-, EU-, and EMS-sEVs demonstrating comparable size profiles across local lesion-associated compartments **(Fig. S1a, b).** In contrast, particle concentration varied significantly by both sample type and disease stage. Plasma from patients with severe EM exhibited significantly higher particle concentrations compared with mild EM and healthy controls (**Fig. 1a**), indicating increased circulating vesicle abundance in severe disease. EM patient tissue-derived sEV preparations from EMS and EU exhibited significantly higher particle concentrations than PF-derived sEV preparations (**Fig. 1b**), suggesting tissue compartment-specific differences in sEV abundance within the EM microenvironment. TEM analysis confirmed the presence of vesicles consistent with standard sEV morphology across all sample types, with minimal background contamination (**Fig. 1c-h**). While all groups exhibited canonical vesicle morphology, qualitative differences in structural heterogeneity were observed between compartments and disease stages. Specifically, severe-stage PF- and EMS-derived sEV preparations demonstrated increased morphological variability relative to other sample groups, with some vesicles exhibiting visible intravesicular electron-dense structures, suggesting differences in sEV ultrastructural features associated with disease stage (**Fig. 1c, f**).

**Figure 1.**
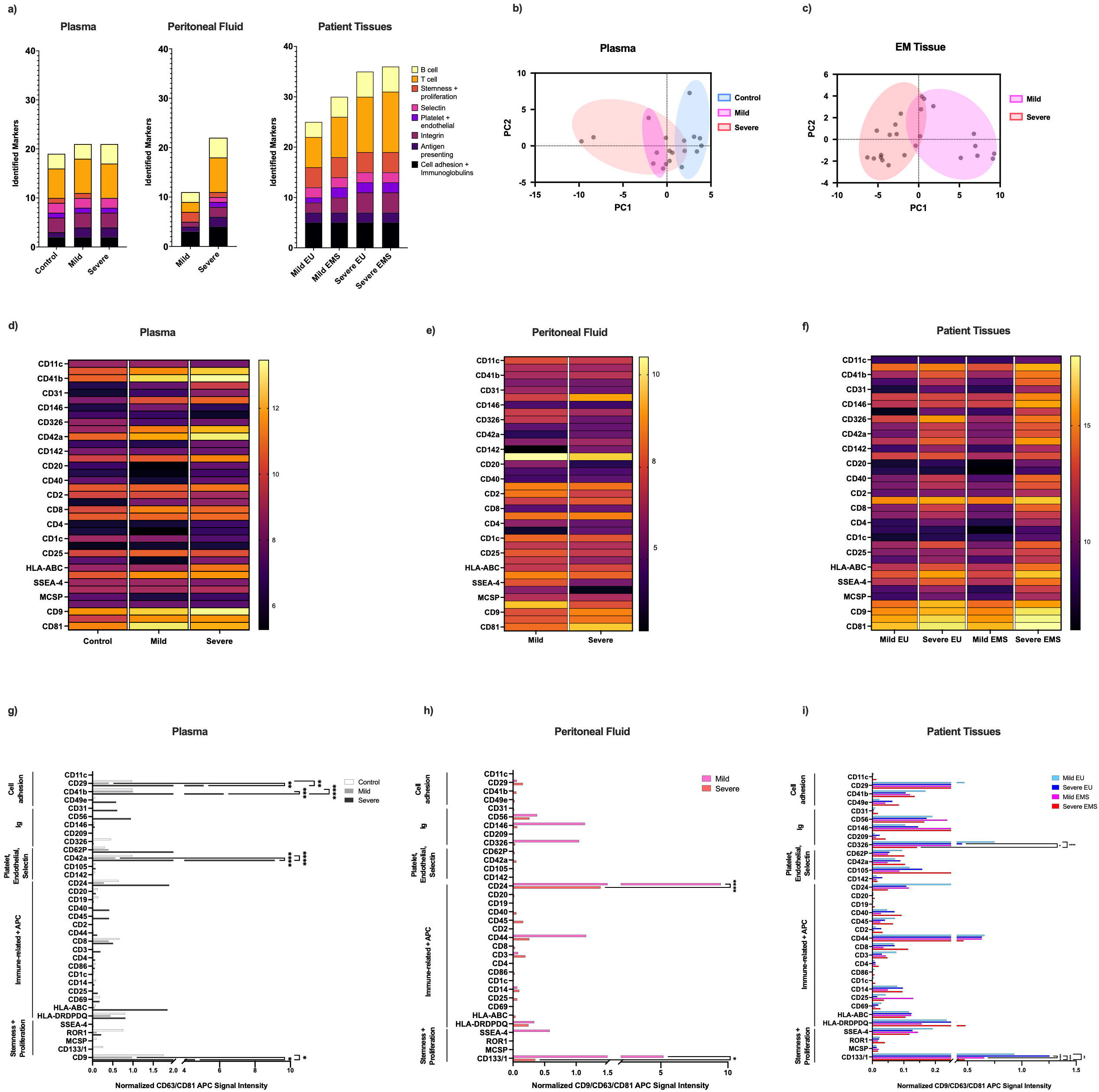
Characterization and surface marker profiling of EM-derived sEVs reveal stage- and tissue-specific differences in vesicle abundance, morphology, tetraspanin expression, and immune- and adhesion-related phenotypes. sEVs were isolated by SEC from matched EMS, EU, PF, and plasma samples from patients with mild-stage EM (n = 5), severe-stage EM (n = 7), as well as plasma samples from healthy controls (n = 8), and assessed for particle concentration, size distribution, morphology, and surface marker composition. **(A-B)** NTA confirmed the presence of abundant sEVs within the expected size range of 50-250 nm across all patient and healthy control samples. **(A)** Median particle concentration per mL of plasma-derived sEVs from patients with mild EM, severe EM, and healthy controls. **(B)** Median particle concentration per mL of matched EMS-, EU-, and PF-derived sEVs from EM patients. (C-H) TEM imaging confirmed die presence of characteristic vesicular structures consistent with sEV morphology across all sample types. Representative TEM images are shown for **(C)** EU-derived sEVs (n = 3), **(D)** EMS-derived sEVs (n = 3), (E-F) PF-derived sEVs (n = 4), **(G)** plasma-derived sEVs from healthy controls (n = 4), and **(H)** plasma-derived sEVs from EM patients (n = 6). **(I-K)** Surface tetraspanin expression (CD9, CD63, and CD81) was assessed using bead-based flow cytometry in isolated sEVs and normalized to bead-only controls in (I) plasma-derived sEVs, (J) PF-derived sEVs, and (K) tissue-derived sEVs. **(L)** MACSPlex EV surface marker profiling identified the proportion of detected surface epitopes across plasma-, PF-, EU-, and EMS-derived sEVs from mild-stage EM patients (n = 5), severe-stage EM patients (n = 7), and healthy controls (n = 8). (M-N) PCAof sEV surface marker profiles across **(M)** patient and control plasma and **(N)** EU- and EMS-derived sEVs from mild- and severe-stage EM patients. **(O-Q)** Expression of selected surface epitopes was normalized to MFI of APC-conjugated tetraspanin signals (CD63/CD81 in panel O; CD9/CD63/CD81 in panels P and Q). Statistical analyses were performed using one-way ANOVA (panels A and B), whereas all other comparisons were performed using two-way ANOVA with Tukey’s multiple comparisons test. *P < 0.05, **P <0.01, ****P < 0.0001. Images were acquired at 94,000* magnification. Scale bars represent 200 nm.

To further characterize sEV phenotype, canonical tetraspanin markers CD9, CD63, and CD81 were assessed using the MACSPlex EV Kit, a bead-based flow cytometry platform (**Fig. 1i-k**). Tetraspanin expression was detected across all sample groups; however, distinct tissue- and stage-associated differences were observed. Namely, CD9 expression was significantly increased in plasma-sEVs from severe EM patients compared with mild EM and control groups (**Fig. 1i**), whereas no significant differences in tetraspanin abundance were detected among PF-sEVs (**Fig. 1j**). In contrast, CD63 and CD81 abundance were increased in severe-stage tissue-sEVs from both EU and EMS compared with mild-stage samples (**Fig. 1k**), suggesting stage-associated remodeling of tissue-sEV populations.

Beyond canonical tetraspanin markers, the MACSPlex analysis was used to assess an additional 37 EV-associated surface epitopes across matched patient samples and control plasma (**Fig. 1l-q**). To visualize broader patterns of sEV surface composition across biological compartments and disease stages, detected markers were grouped into biological categories, including adhesion-associated, immune-associated, antigen-presentation, epithelial-associated, stemness-associated, and platelet-associated markers (**Fig. 1l**). Source-specific heatmap analyses further illustrated compartment- and disease-associated variation in surface marker expression across plasma-, PF-, and tissue-derived sEVs **(Fig. S1c-e)**. Together, these analyses demonstrated distinct compartment-specific patterns of sEV surface composition across sample sources.

Principal component analysis (PCA) demonstrated partial separation of sEV surface marker profiles according to disease stage in plasma-sEVs (**Fig. 1m**) and EU- and EMS-sEVs from mild- and severe-stage EM patients (**Fig. 1n**), indicating disease-associated variation in overall sEV surface composition. At the individual marker level, severe-stage plasma-sEVs demonstrated significantly elevated expression of adhesion- and platelet-associated markers (CD29, CD41b, CD42a, and CD62P) relative to healthy controls, consistent with altered vascular- and platelet-associated signaling in severe disease (**Fig. 1o**). PF-sEVs from mild-stage disease exhibited increased expression of stemness-associated (CD133/1) and immune-modulatory (CD24) compared with severe-stage disease (**Fig. 1p**). In tissue-sEVs, CD133/1 and CD326/EPCAM expression was reduced in EMS compared with matched EU, indicating differences in stemness- and epithelial-associated sEV surface profiles (**Fig. 1q**). Moreover, when comparing expression across disease stages, HLA-II expression was increased in severe-stage tissues, which is consistent with enhanced immune activation in severe lesions.

### 3.2 Proteomic profiling of lesion microenvironment-derived sEVs reveals stage-associated remodeling in EM

Given the disease stage- and tissue-specific differences observed in sEV surface marker profiles, we next investigated whether these phenotypic alterations were accompanied by broader changes in sEV-associated protein cargo. Proteomic profiling of sEVs isolated from matched EU, EMS, and PF revealed pronounced stage- and tissue-dependent remodeling of the sEV proteome across the local EM microenvironment. PCA demonstrated clear separation between mild- and severe-stage tissue-derived sEV samples, consistent with stage-associated proteomic signatures (**Fig. 2a**). Differential protein overlap analyses further supported this divergence, with severe-EMS-sEVs exhibiting substantial expansion of uniquely detected proteins (1012, 16% unique analytes) relative to mild-stage disease. In contrast, EU- and EMS-sEVs within each disease stage shared the majority of detected proteins and exhibited minimal unique representation (<2% unique analytes; **Fig. 2b-d**), indicating that proteomic differences were driven predominantly by disease stage rather than tissue source.

**Figure 2.**
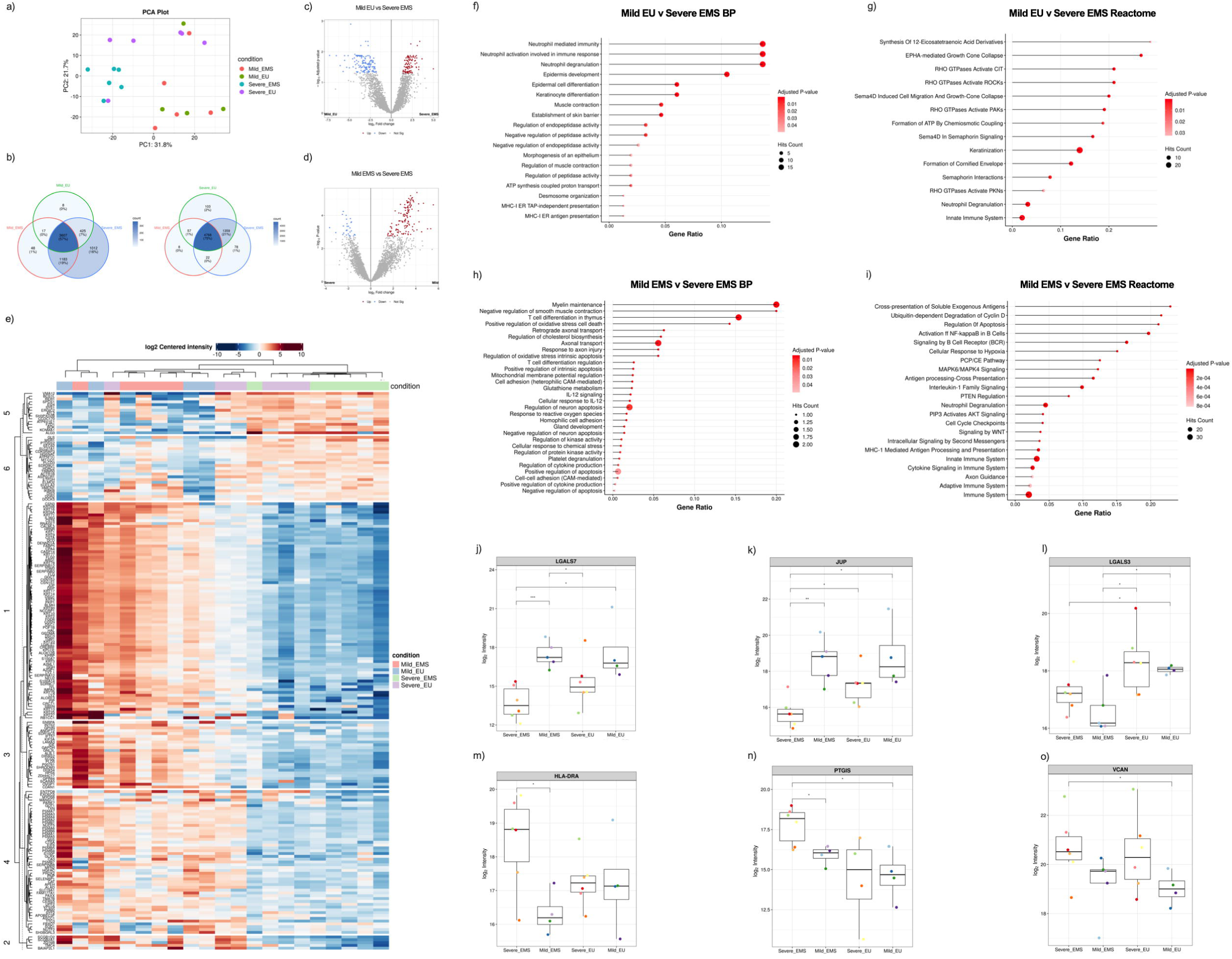
Proteomic profiling of tissue-derived sEV s identifies stage- and tissue-associated differences in protein expression and pathway enrichment in EM. **(A)** PCA of **sEV** proteomes from EU and EMS samples from patients with mild- and severe-stage EM. **(B)** Venn diagrams showing shared and unique proteins detected across mild- and severe-stage tissue-derived sEVs. **(C, D)** Volcano plot analysis identified significantly differentially expressed proteins between mild- and severe-stage tissue-derived sEVs. **(E)** Unsupervised hierarchical clustering heatmap of tissue-derived sEV proteomes. **(F-G)** Functional enrichment analyses of differentially expressed proteins. **(F)** Gene Ontology (GO) Biological Process (BP) enrichment analysis of mild EU versus severe EMS. **(G)** Reactome pathway enrichment analysis of mild EU versus severe EMS. **(H)** GO BP enrichment analysis of mild EMS versus severe EMS. **(I)** Reactome pathway enrichment analysis of mild EMS versus severe EMS. Functional enrichment analyses were performed using over-representation analysis (ORA) with an adjusted p-value cutoff of **0.05. (J-O)** Box plots showing the expression of representative differentially expressed proteins associated with epithelial organization **(J)** LGALS7 and **(K)** JUP, immune modulation **(L)** LGALS3 and **(M)** HLA-DRA, and ECM remodeling (N) PTGIS and **(O)** VCAN. Differential expression analysis was performed using a Benjamini-Hochberg adjusted p-value cutoff of **0.05** and a |log 2 fold-changej > **0.5** following variance-stabilizing normalization (VSN), without data imputation.

Consistent with these findings, unsupervised hierarchical clustering segregated mild- and severe-stage tissue samples into distinct clusters (**Fig. 2e**). Although, one severe-stage EU sample clustered more closely with mild-stage tissues, suggesting partial retention of eutopic-like molecular characteristics in select severe-stage cases. A similar, though less pronounced, pattern was observed within the peritoneal microenvironment, where PF-derived sEV proteomes demonstrated partial stage-associated clustering alongside increased protein diversity in severe-stage disease **(Fig. S2a-c)**, consistent with remodeling of the sEV proteome during EM progression.

To define the biological programs underlying these stage-associated proteomic shifts, pathway enrichment analyses were performed. Comparisons between mild-stage EU and severe-stage EMS demonstrated broad enrichment of inflammatory and immune-associated signaling networks, including cytokine signaling, antigen presentation, innate immune activation, cellular stress responses, and apoptosis-related pathways (**Fig. 2f, g**), consistent with establishment of an increasingly immune-active lesion microenvironment in advanced disease. In parallel, comparisons between mild- and severe-EMS-sEV proteomes identified enrichment of pathways linked to epithelial remodeling, cytoskeletal organization, adhesion dynamics, and immune regulation (**Fig. 2h, i**), supporting progressive restructuring and influence on shaping the dynamic lesion microenvironment during disease progression. At the protein level, these alterations reflected coordinated shifts between epithelial maintenance-associated and inflammatory remodeling-associated programs. Mild-stage disease was characterized by enrichment of proteins linked to epithelial integrity and junctional stability, including galectin-7 (LGALS7) and junction plakoglobin (JUP), which were preferentially enriched in mild-stage EMS and EU tissues relative to severe-stage samples (**Fig. 2j, k**). In contrast, severe-stage disease was characterized by enrichment of proteins associated with immune modulation, including galectin-3 (LGALS3) and HLA-DRA, as well as ECM remodeling, including prostacyclin synthase (PTGIS) and versican (VCAN) (**Fig. 2l-o**).

Consistent with tissue-derived findings, PF-derived sEV proteomes demonstrated stage-associated alterations linked to cell adhesion and tissue remodeling in mild-stage disease and altered metabolic and ECM-associated processes in severe-stage disease **(Fig. S2d-k)**. Biological process and pathway enrichment analyses revealed that severe-stage PF-derived sEVs were enriched for proteins associated with membrane remodeling, ECM biosynthesis, metabolic reprogramming, and inflammatory stress adaptation, consistent with progression toward a chronic inflammatory and lesion permissive state **(Fig. S2d, e).** Notably, several proteins demonstrated compartment-specific enrichment patterns between tissue- and PF-derived sEVs, suggesting selective extracellular distribution of stage-associated signaling programs within the lesion microenvironment **(Fig. S3).** Proteins linked to remodeling (PROM1 or CD133, MGAT1) and proliferative-associated processes (MYOF, TGFBR3) were preferentially enriched in mild-stage PF-derived sEVs despite reduced abundance in corresponding tissue-derived populations **(Fig. S3e,f)**, whereas proteins associated with inflammatory stress adaptation and proteostatic regulation (XRCC5, PSMD9) demonstrated concordant enrichment across severe-PF- and tissue-sEVs **(Fig. S3g)**. PF-derived sEV proteomes demonstrated partial overlap with tissue-derived stage-associated signatures, while also exhibiting distinct enrichment patterns associated with the peritoneal microenvironment. Collectively, these findings identify coordinated but compartment-specific stage-associated sEV proteomic programs in EM.

### 3.3 Plasma-derived sEV proteomes reflect progressive systemic immune, vascular and ECM-remodeling programs in EM

To determine whether the stage-associated molecular remodeling observed within lesion compartments was reflected systemically, we performed proteomic profiling of plasma-sEVs from EM patients and healthy controls. PCA demonstrated clustering of EM patient samples relative to healthy controls. In contrast to the stage-associated separation observed in tissue-sEV proteomes (**Fig. 2a**), plasma-derived sEVs exhibited less distinct separation between mild- and severe-stage disease, consistent with a more conserved systemic disease-associated sEV signature across EM stages (**Fig. 3a**). Differential protein overlap analyses identified both shared and stage-specific systemic signatures, including a subset of uniquely detected proteins in severe-stage plasma-sEVs (117, 9% unique analytes), consistent with progressive systemic proteomic remodeling in severe disease (**Fig. 3b, c**). Unsupervised hierarchical clustering largely separated patient and control plasma samples. Though, partial clustering of mild-stage patients with controls suggested that mild disease retains a more homeostatic systemic sEV profile, which becomes lost in a stage-dependent manner (**Fig. 3d**).

**Figure 3.**
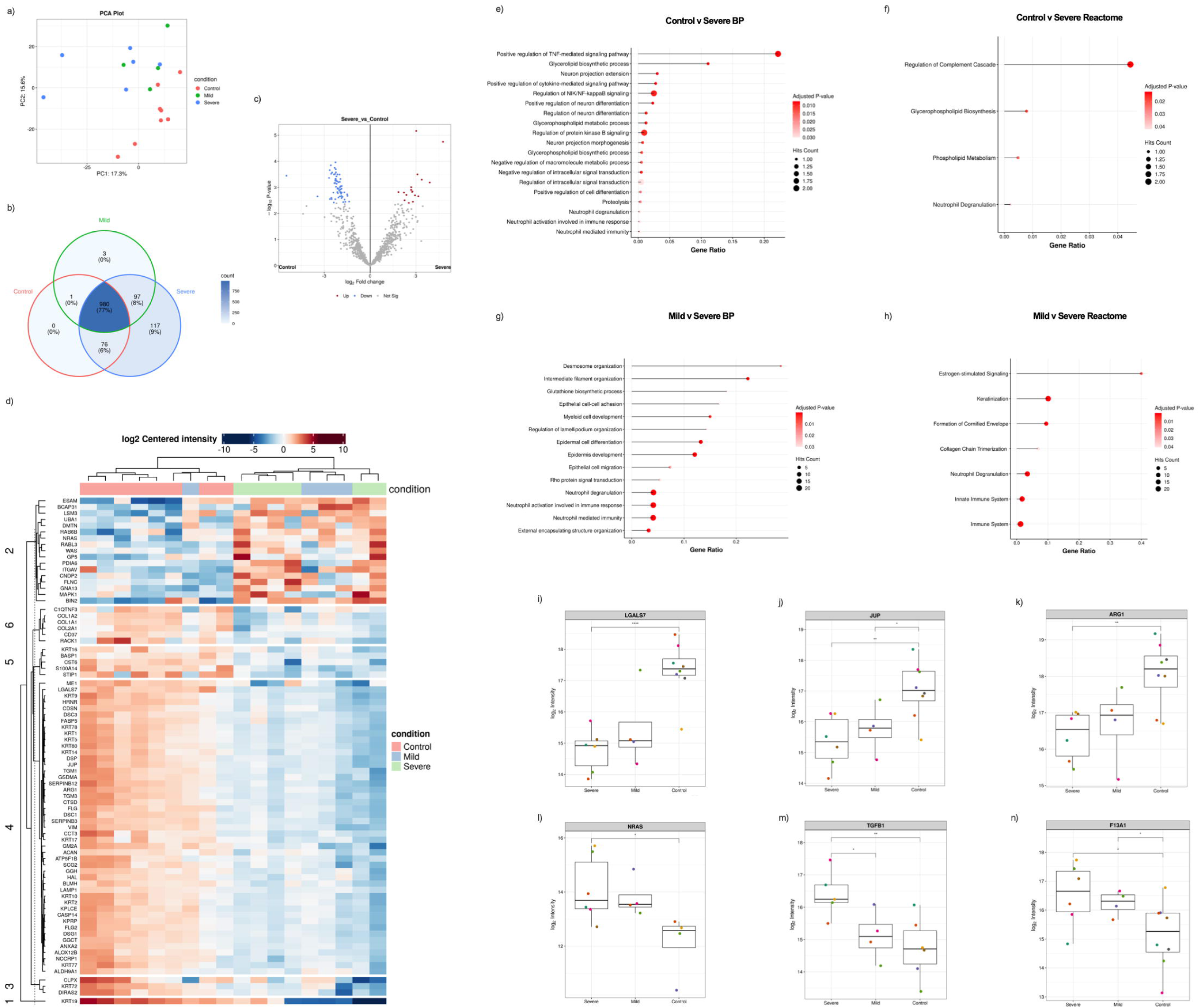
Proteomic profiling of plasma-derived sEVs identifies disease stage-associated differences in protein expression and pathway enrichment in EM. **(A)** PCAof plasma-derived sEV proteomes from patients with mild- and severe-stage EM and healthy controls. **(B)** Venn diagram showing shared and unique proteins detected across healthy control, mild-stage, and severe-stage plasma-derived sEVs. **(C)** Volcano plot analysis identified significantly differentially expressed proteins between severe-stage and control plasma-derived sEV **(D)** Unsupervised hierarchical clustering heatmap of plasma-derived sEV proteomes. **(E-H)** Functional enrichment analyses of differentially expressed proteins from plasma-derived sEVs from plasma-derived sEVs. (E, F GO Biological Process and Reactome pathway enrichment analyses comparing severe-stage patients with controls. **(G, H)** GO Biological Process and Reactome pathway enrichment analyses comparing mild- and severe-stage patient-derived sEVs. **(I-N)** Box plots showing die expression of representative differentially expressed plasma-derived sEV proteins associated with epithelial organization **(I)** LGALS7 and **(J)** JUP, immune regulation **(K)** ARG1, and vascular remodeling and pro-fibrotic signaling **(L)** NRAS, **(M)** TGFB1, and **(N)** F13A1. Differential expression analysis was performed using a Benjamini-Hochberg adjusted p-value cutoff of 0.05 and log2 fold-change threshold 2:0.5 following variance stabilizing normalization (VSN), without data imputation.

To define the biological programs underlying systemic sEV remodeling, pathway enrichment analyses were performed. Comparisons between mild-stage patients and healthy controls revealed enrichment of inflammatory signaling and membrane remodeling pathways, including innate immune activation, cytokine-associated signaling, complement activation, and lipid metabolic processes, consistent with systemic immune activation and membrane lipid remodeling in mild EM (**Fig. 3e, f**). In contrast, severe plasma-sEVs demonstrated enrichment of epithelial remodeling programs coupled with stress- and inflammation-associated signaling pathways, including Rho GTPase-linked cytoskeletal regulation and keratinization-associated processes (**Fig. 3g, h**).

At the protein level, systemic alterations reflected coordinated changes in epithelial integrity, immune regulation, and remodeling-associated signaling. Proteins associated with epithelial maintenance and junctional stability, including LGALS7 and JUP, were preferentially enriched in control plasma-sEVs relative to both mild- and severe-stage patient samples (**Fig. 3i, j**), consistent with a loss of epithelial homeostasis in severe disease. Arginase-1 (ARG1) was similarly elevated in controls, supporting altered systemic immune regulatory and myeloid-associated signaling during EM progression (**Fig. 3k**). In contrast, severe plasma-sEVs demonstrated enrichment of proteins associated with inflammatory persistence, vascular remodeling, and pro-fibrotic signaling, including NRAS, TGFB1, and coagulation factor XIII A chain (F13A1) (**Fig. 3l-n**), consistent with enhanced systemic inflammatory signaling, vascular activation, and ECM stabilization in severe disease.

Comparisons of plasma- and tissue-derived sEV proteomes revealed both conserved and compartment-specific signatures. Proteins associated with epithelial adhesion (DSC1, PKP1) and homeostasis (CSTA, POF1B) were enriched in control plasma and mild-stage tissues **(Fig. S3a)**, whereas proteins linked to inflammatory and remodeling-associated programs (TGFB1, FN1, ICAM1, and F13A1) demonstrated concordant enrichment across severe-stage plasma and EMS-derived sEVs **(Fig. S3b).** These findings support partial reflection of lesion-associated remodeling programs within the systemic circulation. Immune-associated proteins displayed both shared and compartment-specific regulation across disease states, with antigen-presentation and inflammatory activation markers enriched in severe-stage tissues and plasma-derived sEVs (HLA-related proteins, B2M) **(Fig. S3c)**. Proteins associated with immune homeostasis and neutrophil-associated responses were relatively enriched in healthy controls (CD37, IL36G), suggesting systemic immune remodeling during disease progression **(Fig. S3c)**. Additionally, proteins associated with immune modulation and tissue adaptation (ATG1 and S100A7) were enriched in mild-stage EMS-sEVs relative to severe-stage disease samples, potentially supporting a role for these processes in lesion development **(Fig. S3d).**

Collectively, these findings support the emergence of coordinated systemic and lesion-associated sEV proteomic programs in EM.

### 3.4 Lipidomic profiling of lesion-derived sEVs reveals stage-associated remodeling in EM

While proteomic and surface marker profiling provide important insight into the molecular cargo and immunological landscape of sEVs, they do not fully capture alterations in membrane lipid composition that accompany vesicle structure and function. As major structural components of sEVs, lipids regulate membrane organization, cargo packaging, and recipient cell interactions. Although metabolic dysregulation is increasingly recognized as an important feature of EM pathophysiology, the lipid composition of disease-associated sEVs remains poorly understood. We therefore incorporated lipidomic profiling to identify stage- and tissue-specific metabolic signatures and to complement proteomic characterization of EM-derived sEVs.

Lipidomic profiling of tissue-derived sEVs from a separate patient cohort containing pooled EU samples (n = 4; combined mild- and severe-stage due to limited sample availability) and EMS from patients with mild (n = 4) and severe (n = 4) EM revealed pronounced stage- and tissue-specific alterations in lipid composition, predominantly in EMS-derived sEVs. Lipid species were analyzed using LC-MS in both positive and negative electrospray ionization modes to improve lipidome coverage, as complementary ionization strategies enable detection of distinct subsets of lipid species based on their physical and chemical properties. PCA revealed clear separation between EU and severe-stage EMS-sEVs, and most notably between mild- and severe-EMS-sEVs (**Fig. 4a, d, j, m**). Differentially expressed lipids were visualized in an unsupervised heatmap, revealing distinct clustering patterns among disease groups based on their lipid composition (**Fig. 4b, e, k, n**). To further define the biological programs underlying these lipidomic alterations, metaboanalyst enrichment was performed.

**Figure 4.**
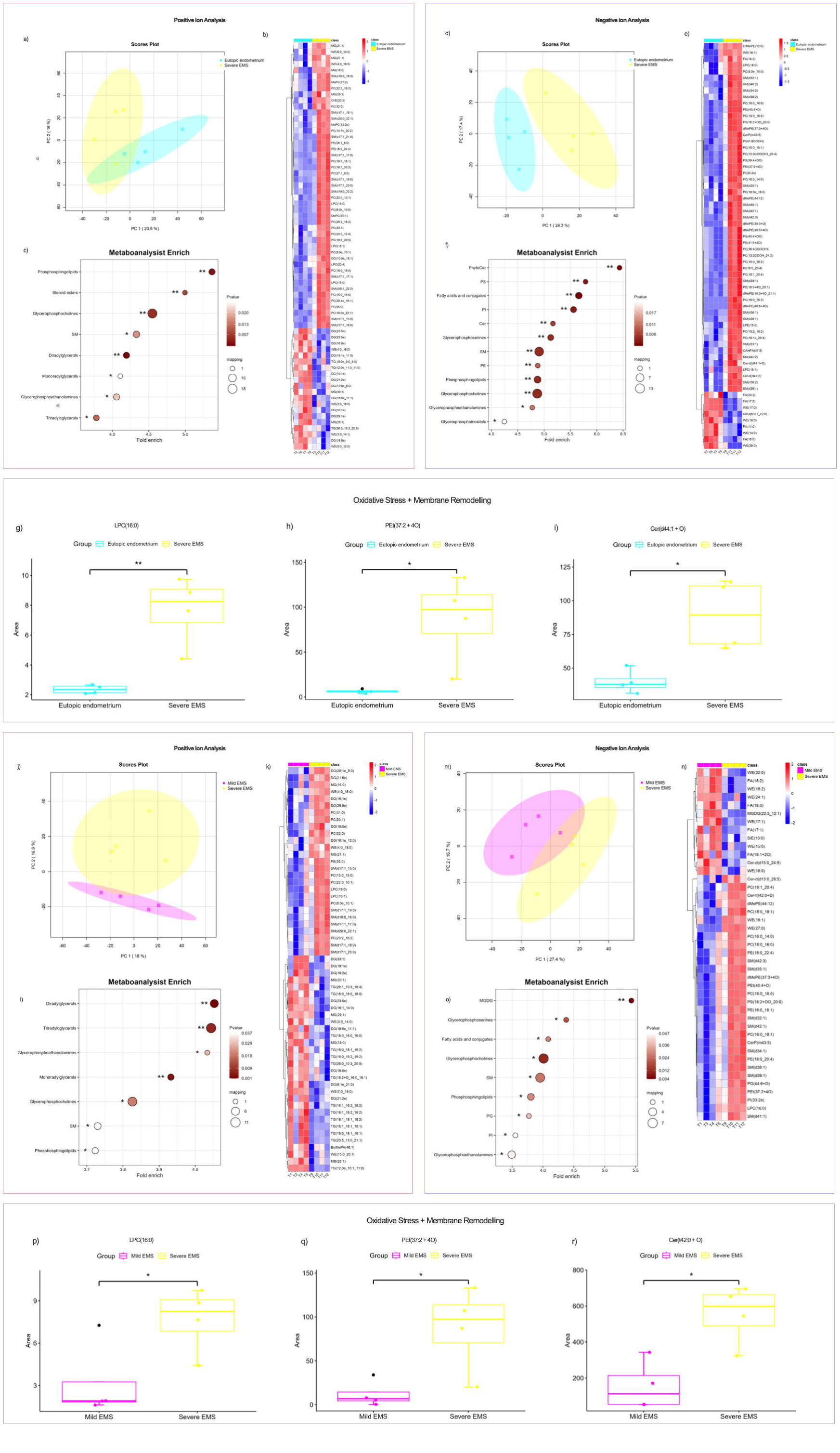
Lipidomic profiling of tissue-derived sEVs identifies differential lipid signatures across tissue compartments and disease stages. Lipid composition of tissue-derived sEVs from EU (n = 4), mild-EMS (n = 4), and severe-EMS; n=4) was assessed by LC-MS using **(A-C, J-L)** positive and **(D-F, M-O)** negative electrospray ionization modes **(A)** PCA of lipid profiles from severe-EMS and EU sEVs. **(B)** Heatmap analysis of lipid species detected in severe-EMS and EU sEVs. **(C)** Dot plot enrichment analysis showing lipid class representation in severe-EMS vs EU sEVs samples. **(D)** PCA of lipid profiles from severe-EMS and EU sEVs. **(E)** Heatmap analysis of lipid species detected in severe-EMS and EU sEVs. **(F)** Dot plot enrichment analysis showing lipid class representation in severe-EMS vs EU sEVs samples. **(G-I)** Representative box plots show relative abundance of select lipid species in severe-EMS compared with EU sEVs, including **(G)** LPC(16:0), **(H)** PEt(37:2 + 40), and **(I)** Cer(d44:l + O). **(J)** PCA of lipid profiles from mild- and severe-EMS sEVs. **(K)** Heatmap analysis of lipid species detected in mild- and severe-EMS sEVs. **(L)** Dot plot enrichment analysis showing lipid class representation in severe-EMS vs mild-EMS sEV samples. **(M)** PCA of lipid profiles from mild- and severe-EMS sEVs. **(N)** Heatmap analysis of lipid species detected in mild- and severe-EMS sEVs. **(O)** Dot plot enrichment analysis showing lipid class representation in severe-EMS vs mild-EMS samples. **(P-R)** Representative box plots show select lipid species in severe-EMS compared with mild-stage EMS sEVs, including **(P)** LPC(16:0), **(Q)** PEt(37:2 + 40), and **(R)** Cer(d42:0 + O). Analysis was filtered and confirmed by combining the results of the VIP values (VIP >1.5), fold-change (|log2 FC| > 1) and t-test (*P < 0.05).

Comparisons between severe-EMS and EU sEVs demonstrated enrichment of multiple membrane-associated lipid classes, including sphingomyelins (SM), glycerophosphocholines (GPC), phosphatidylethanolamines (PE), phosphatidylserines (PS), phosphatidylinositols (PI), and ceramides (Cer), together with increased abundance of neutral glycerolipids (DG, MAG, and TG; **Fig. 4c, f**). Similarly, comparisons between mild- and severe-EMS-sEVs identified enrichment of membrane phospholipids and sphingolipids alongside neutral lipid classes, including SM, PI, PG, DG, MAG, TG, and monogalactosyldiacylglycerols (MGDG) in severe-EMS, supporting membrane remodeling and altered lipid metabolism associated with severe EM (**Fig. 4l, o**).

At the individual lipid level, severe-EMS-sEVs demonstrated increased abundance of lysophosphatidylcholine (LPC 16:0), oxidized phosphatidylethanolamine (PEt 37:2 + 4O), and oxidized ceramide species (Cer d44:1 + O, Cer t42:0 + O) relative to both EU samples and mild-stage EMS (**Fig. 4g-i, p-r)**. These alterations are consistent with coordinated remodeling of membrane architecture together with increased oxidative lipid signaling in severe disease. In contrast, comparisons between mild-EMS- and EU-sEVs demonstrated modest lipidomic remodeling **(Fig. S4)**. Mild-stage lesions were enriched in neutral lipid classes, including MAG, DG, and TG, together with select fatty acid, MGDG, Cer, and hexosylceramide (HexCer) species. These findings suggest that mild-EMS-sEVs likely undergo selective membrane remodeling, although to a lesser extent than the oxidative and membrane-associated lipid alterations observed in severe disease.

### 3.5 Plasma-derived sEV lipidomic profiling identifies disease-associated systemic lipid remodeling in EM

Having identified disease stage-dependent remodeling of EMS-derived sEV lipidome, we next examined whether circulating plasma-sEVs from a separate patient cohort containing mild-stage (n=6), severe-stage (n=6) and healthy controls (n=8), would reveal disease-specific enrichment. Indeed, PCA and hierarchical clustering analyses clearly demonstrated separation between severe-stage EM and control plasma samples, indicating substantial systemic remodeling of the circulating sEV lipidome (**Fig. 5a, b, g, h**). Metaboanalyst enrichment revealed increased representation of multiple membrane-associated lipid classes, including GPC, PE, SM, Cer, and neutral glycerolipids (TG and MAG), consistent with widespread remodeling of circulating membrane lipids during advanced disease (**Fig. 5c, i**). At the individual lipid level, severe plasma-sEVs demonstrated increased abundance of oxidized triglycerides (TG 54:5 + O), oxidized phosphatidylcholines (PC 36:6 + OO), lysophosphatidylcholines (LPC 16:0), and glycosphingolipid species (Hex2Cer d30:0, Hex2Cer d29:0 + 2O) together with reduced CoQ10, collectively supporting increased oxidative stress, altered membrane remodeling, and mitochondrial dysfunction in severe EM (**Fig. 5d-l**). Comparisons between mild-EM and controls similarly demonstrated distinct lipidomic differences, with PCA, heatmap visualization, and lipid class enrichment analyses revealing separation between groups and altered lipid composition patterns **(Fig. S4)**, indicating that systemic alterations in the circulating sEV lipidome are detectable in mild-stage disease.

**Figure 5.**
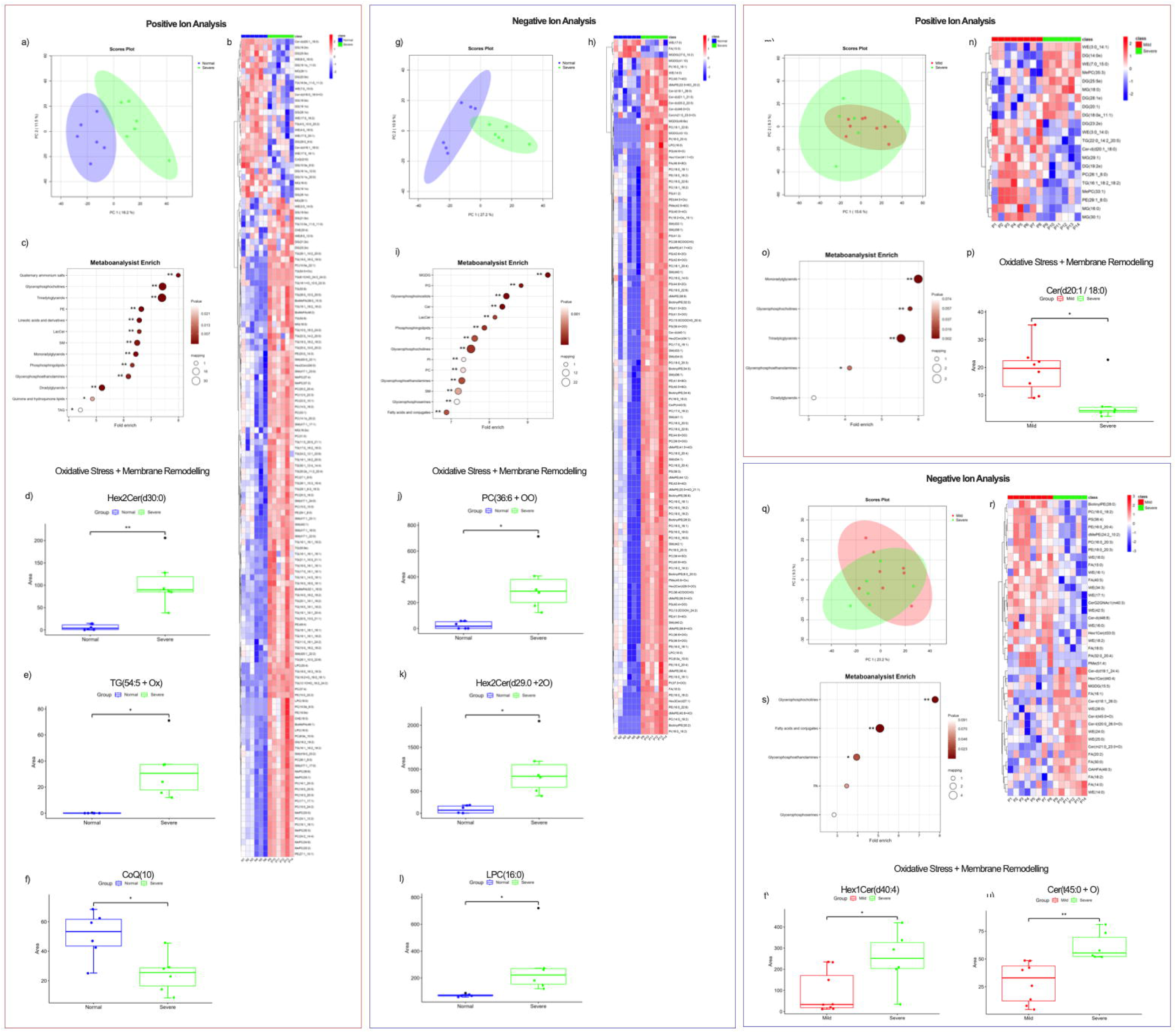
Plasma-derived sEV lipid profiles exhibit coordinated phospholipid remodeling and systemic oxidative stress-associated lipid signatures in EM. Lipid composition of tissue-derived sEVs fiom EU (n = 4), mild-EMS (n=4), and sevcro-EMS; n=4) was assessed by LC-MS using (A-F, M-P) positive and (G-L, Q-LJ) negative electrospray ionization modes (A) PCA demonstrated separation between severe-stage and control plasma sEV-dcrived lipid profiles, indicative of systemic lipid remodeling **(B)** Heatmap analysis revealed clear separation between severe-stage and control plasma samples, supporting lipidomic divergence associated with disease presence. (C) Dot plot enrichment analysis demonstrated over-representation of GPC, TG, PE, *Cer,* SM and MAG in scvcrc-plasma samples. (D-F) Representative box plots demonstrated that scvcre-plasma sEVs exhibited increased abundance of (D) Hex2Cer(d30:0), **(E)** TG(54:5 + O), alongside a significant decrease in (F) CoQ( 10) relative to control plasma. (G) PCA demonstrated separation between severe-stage and control plasma sEV-derivcd lipid profiles **(H)** Heatmap analysis revealed clear separation between severe-stage and control plasma samples. **(I)** Dot plot enrichment analysis demonstrated over-representation of MCDG, PC, PG, Cer, SM, PS and GPC in scvcre-plasma samples. (J-L) Representative box plots demonstrated that sevcrc-plasma exhibited increased abundance of (J) PC(36:6+OO), **(K)** Hex2Cer(d29:0+20), and (L) LPC(16:0), relative to control plasma. (M) PCA demonstrated overlap between mild- and severe-stage plasma sEV-dcrivcd lipid profiles. **(N)** Heatmap analysis revealed partial separation between mild- and scvcrc-plasma samples, supporting lipidomic remodeling associated with disease stage. **(O)** Dot plot enrichment analysis demonstrated over-representation of MG, GPC, TG and DG in scvcrc-plasma samples. (P) Representative box plots demonstrated that mild-plasma samples exhibited increased abundance of Cer(d20:1 /18:0) (Q) PCA demonstrated overlap between mild- and severe plasma sEV-derived lipid profiles. **(R)** Heatmap analysis revealed partial separation between mild- and scvcrc-plasma samples. (S) Dot plot enrichment analysis demonstrated over-representation of GPC, FA, PA, and GPS in scvcrc-plasma samples. **(T-U)** Representative box plots demonstrated that scvcrc-plasma exhibited increased abundance of **(T)** HexlCeifd40:4) and **(U)** Cer(t45:0 + O) relative to mild plasma. Analysis was filtered and continued by combining the results ofthe VIPvalues(VIP>1.5), fold-change (|log2FC| > 1) and t-test(*P< 0.05, **P<0.01).

Direct comparison of mild- and severe-plasma-sEVs demonstrated comparatively modest stage-associated differences, with partial overlap observed by PCA despite selective enrichment of glycerophospholipid (GP), fatty acid (FA), and ceramide (Hex1Cer(d40:4), Cer(t45:0 + O)) classes in severe disease (**Fig. 5m-u**). Plasma-sEVs demonstrated fewer stage-dependent differences compared with EMS-sEVs. Several lipid species, including LPC(16:0), oxidized triglycerides, and glycosphingolipids, were consistently altered across tissue- and plasma-derived sEV populations.

### 3.6 Integrated proteome and lipidome signatures identify coordinated molecular remodeling of severe-EMS-derived sEVs

Integrated multi-omics analyses were performed to determine whether stage- and tissue-dependent alterations identified across individual proteomic and lipidomic datasets represented coordinated molecular programs. Following preprocessing and integration of proteomic and lipidomic features, intra- and inter-omics correlation analyses demonstrated covariance within and between molecular datasets in both tissue- and plasma-derived sEVs, supporting relationships between complementary molecular cargo classes **(Fig. S5a, d; S6a, d)**. Proteome-lipidome covariance was greatest within lesion-derived sEVs (positive ion RV = 0.66), whereas plasma-derived sEVs demonstrated moderate integration across positive (RV = 0.40) and negative ion datasets (RV = 0.63). Unsupervised hierarchical clustering of integrated protein and lipid features (displayed in the upper and lower portions of the heatmaps, respectively) demonstrated stage-associated grouping of tissue- and plasma-sEV samples, with more pronounced separation observed between mild- and severe-tissue-sEVs compared with circulating plasma-sEVs **(Fig. S5b, e; S6b, e)**. Correlation network analyses further identified extensive positive and negative associations between protein and lipid features across both tissue- and plasma-derived sEVs, revealing interconnected molecular relationships between distinct cargo classes **(Fig. S5c, f; S6c, f)**.

Building on these findings, multiple co-inertia analysis (MCIA) was performed to evaluate concordant variation between proteomic and lipidomic profiles at the sample level. Integrated tissue-derived sEV profiles demonstrated separation between severe-EMS and EU samples, while mild-EMS samples exhibited greater similarity to EU-derived profiles, suggesting progressive molecular divergence in severe EM (**Fig. 6a,d**).

**Figure 6.**
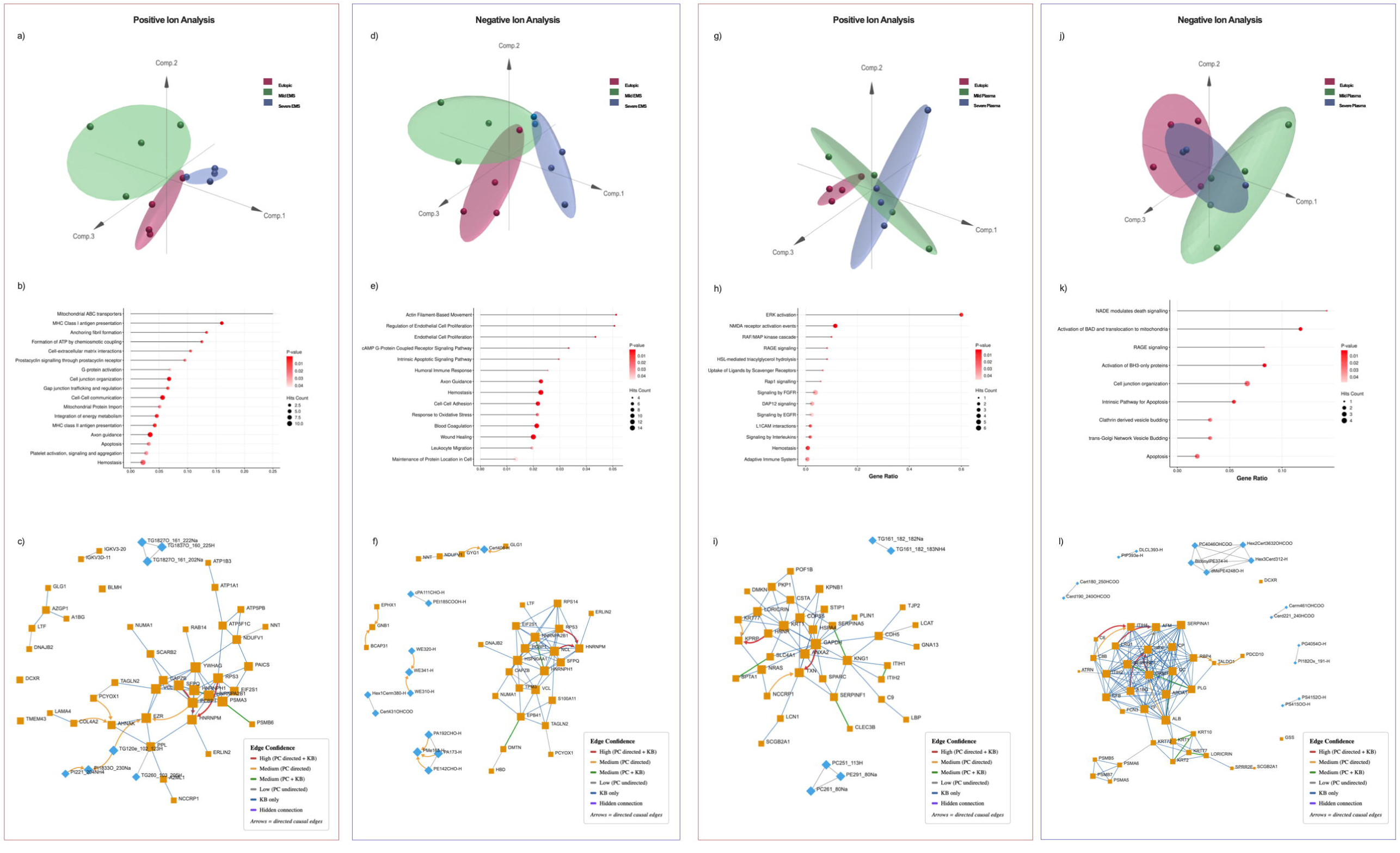
Integrated proteomic and lipidomic analyses identify coordinated molecular signatures associated with severe-EM tissue- and plasma-derived sEVs. (A-C) Positive ion tissue analysis **(A)** MCIA demonstrated separation of severe EMS and EU samples. **(B)** Reactome pathway enrichment identified metabolic, membrane-associated, immune, and cellular organization pathways. **(C)** Causal discovery analysis identified mitochondrial bioenergetics, membrane remodeling, cellular stress, and cytoskeletal organization networks. **(D-F)** Negative ion tissue analysis **(D)** MCIA demonstrated separation of severe EMS and EU samples. **(E)** Reactome analysis identified metabolic, endothelial-, immune-, and membrane remodeling-associated pathways. **(F**) Causal discovery analysis identified epithelial remodeling, vascular signaling, oxidative stress, and phospholipid/ceramide remodeling networks. **(G-I)** Positive ion plasma analysis **(G)** MCIA demonstrated partial separation of severe- and control plasma-derived sEVs. **(H)** Reactome analysis identified immune and intercellular communication, cell junction and ECM organization, mitochondrial metabolism, and platelet/hemostatic signaling pathways. **(I)** Causal discovery analysis identified phospholipid remodeling, epithelial and ECM organization, innate immune signaling, and oxidative stress networks. **(J-L**) Negative ion plasma analysis **(J)** MCIA demonstrated partial separation of severe- and control plasma-derived sEVs. **(K)** Reactome analysis identified vascular remodeling, endothelial and platelet signaling, immune responses, cell adhesion and migration, oxidative stress, and apoptosis pathways. **(L)** Causal discovery analysis identified innate immune and complement activation, lipid transport, phospholipid/sphingolipid remodeling, epithelial remodeling, proteostasis, and metabolic adaptation networks.

Plasma-derived sEV profiles showed partial separation between severe-plasma and controls, whereas mild-plasma-sEVs displayed greater overlap with control profiles, suggesting progressive systemic molecular remodeling in severe EM that remains less pronounced than the signatures observed in tissue-derived sEVs (**Fig. 6g, j**).

To identify the biological programs associated with the integrated molecular features distinguishing tissue compartments and disease stages, Reactome pathway enrichment analysis was performed. In tissue-derived sEVs, comparisons between severe-EMS and EU samples revealed enrichment of metabolic, membrane-associated, immune, and cellular organization pathways in positive ion mode (**Fig. 6b**), whereas negative ion mode analysis identified additional enrichment of endothelial-, immune-, and membrane remodeling-associated pathways (**Fig. 6e**). In plasma-derived sEVs, integrated analysis of severe-stage patients relative to healthy controls identified enrichment of pathways associated with immune and intercellular communication, cell junction and ECM organization, mitochondrial metabolism, and platelet/hemostatic signaling in positive ion mode (**Fig. 6h**). Negative ion plasma analysis similarly demonstrated enrichment of endothelial-, platelet-associated, immune, oxidative stress-, and apoptosis-related pathways in severe-stage patients (**Fig. 6k**). These coordinated pathway enrichments suggest that the integrated molecular signatures identified by multi-omic analyses reflect interconnected biological processes rather than isolated protein or lipid alterations, prompting further investigation of feature-level molecular relationships.

To further investigate relationships between individual molecular features, causal discovery analysis was performed using integrated proteomic and lipidomic datasets to infer potential regulatory connections within sEV-associated molecular networks. These analyses identified relationships between molecular features that extended beyond individual protein or lipid alterations. Tissue-derived sEV networks revealed interconnected modules associated with mitochondrial bioenergetics, membrane lipid remodeling, cellular stress responses, cytoskeletal organization, epithelial remodeling, vascular-associated signaling, and oxidative stress responses (**Fig. 6c,f**). Plasma-derived sEV networks demonstrated interconnected modules involving phospholipid and sphingolipid remodeling, epithelial and ECM organization, innate immune signaling, oxidative stress, lipid transport, proteostasis, and metabolic adaptation (**Fig. 6i,l**).

Collectively, integrated proteomic and lipidomic analyses reveal coordinated stage- and tissue-associated sEV molecular programs during EM progression. Tissue-derived sEVs demonstrated increasingly distinct lesion-associated molecular signatures involving immune activation, ECM organization, vascular signaling, and metabolic adaptation, whereas plasma-derived sEVs reflected parallel systemic inflammatory, vascular, and metabolic remodeling that became more apparent with advancing disease despite greater overlap between integrated profiles.

### 3.7 EM-derived sEVs induce stage- and tissue-specific functional alterations in human uterine microvascular endothelial cells

To determine whether stage-specific molecular differences in EMS-derived sEVs were translated into functional effects, we assessed their impact on endothelial uptake and angiogenic activity in human uterine microvascular endothelial cells (HUtMECs). Cells were treated with MemGlow™-labeled sEVs derived from mild or severe EMS lesions and monitored by live-cell imaging over a 10 h time course. HUtMEC uptake of both mild- and severe-EMS-sEVs was visualized over time, with minimal intracellular fluorescence immediately following treatment and increased signal observed at 4 h post-incubation (**Fig. 7a,b**). EMS-derived sEV-associated fluorescence overlapped with the cytoplasmic stain BODIPY TR, indicating intracellular accumulation within the cytoplasm following endothelial cell uptake (**Fig. 7c, d**). In contrast, sEV-associated fluorescence did not overlap with Hoechst-labeled nuclei at any time point, suggesting that internalized sEVs remained predominantly excluded from the nuclear compartment (**Fig. 7a, b**).

**Figure 7.**
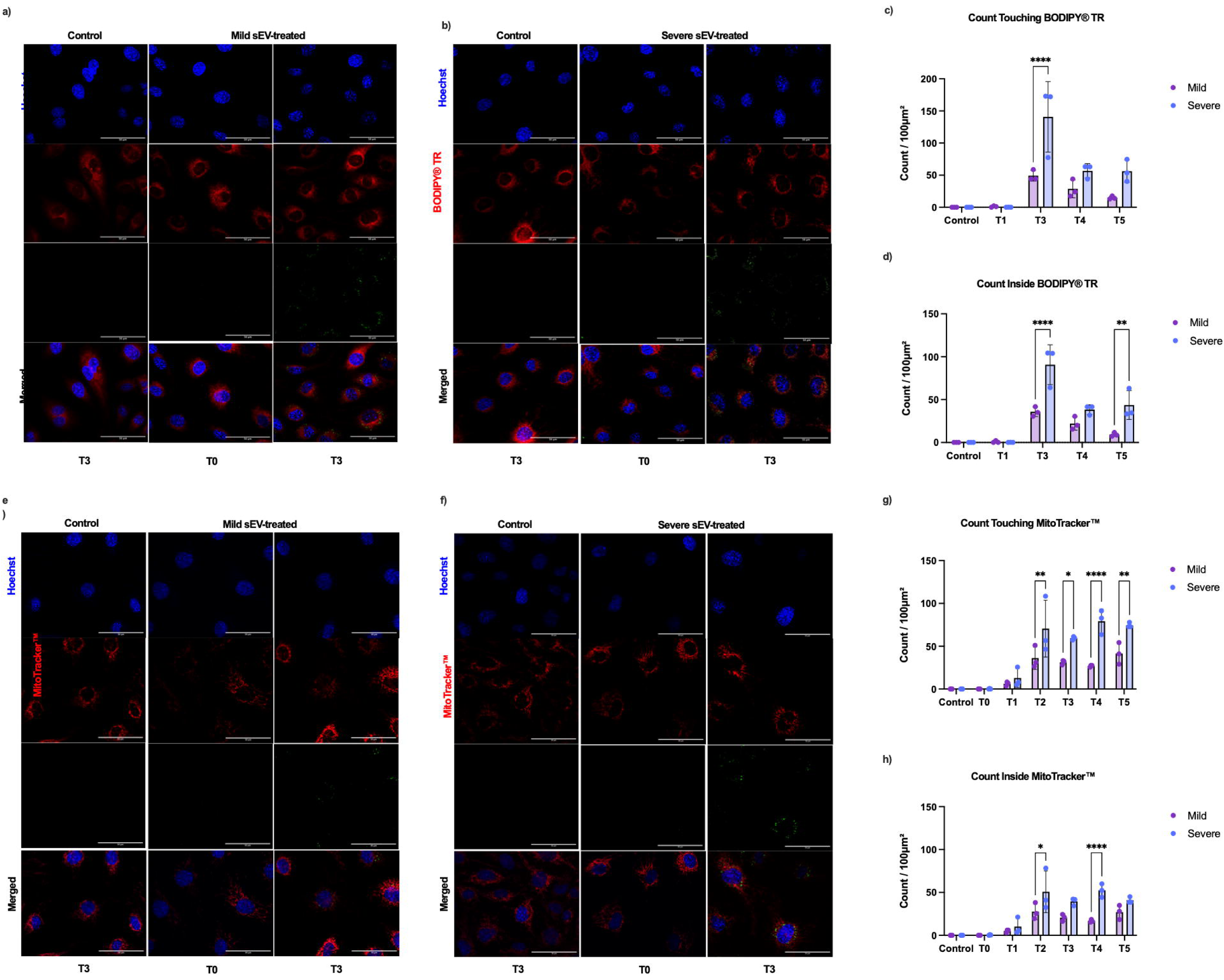
Mild- and severe-EMS-derived sEVs exhibit progressive uptake and mitochondrial localization in HUtMECs. HUtMECs were treated with MemGlow™-labeled pooled mild- or severe-EMS-derived sEVs and imaged by live-cell microscopy over a 10 h time course. **(A, B)** Representative live-cell images showing uptake of MemGlow™-labeled **(A)** mild- and (B) severe-EMS-derived sEVs. Nuclei were stained with Hoechst (blue), the cytoplasm with BODIPY™ TR (red), and sEVs with MemGlow™ (green). Images are shown at the time of plating (TO) and 6 h post-incubation (T3). Unheated control cells received vehicle only (PBS) and exhibited no detectable MemGlow™ fluorescence. **(C, D) Q**uantification of MemGlow™ fluorescence **(C)** associated with and **(D)** internalized by HUtMECs demonstrated progressive accumulation of sEV signal over time (T0-T5,0 −10 h) **(E, F)** Representative live-cell images showing mitochondrial localization of MemGlow™-labeled (E) mild- and (F) severe-EMS-derived sEVs. Nuclei were stained with Hoechst (blue), mitochondria with MitoTracker™ Deep Red (red), and sEVs with MemGlow™ (green). Images are shown at TO and T3. Untreated control cells exhibited no detectable MemGlow™ fluorescence. **(G, H)** Quantification of MemGlow™ fluorescence **(G)** associated with mitochondria and **(H)** the proportion of mitochondrial co-localization demonstrated progressive localization of sEVs to mitochondria over time (T0-T5,0 -lOh). Representative images were acquired using the Leica MICA at 60 magnification. Scale bar = 50 pm. Data are presented as mean ± SD. Statistical significance was determined by two-way ANOVA with appropriate multiple-comparisons testing; *P < 0.05, **P < 0.01, ***P < 0.001, ****p < 0.0001. N = 3 pooled biological replicates.

Following sEV uptake, the intracellular fluorescence pattern observed within HUtMECs demonstrated a morphology resembling mitochondrial structures, suggesting potential mitochondrial association of internalized EMS-sEVs. Given the enrichment of mitochondrial-associated proteins identified through proteomic profiling, (ATP5F1A-C, NDUF6,8,10, COX4I1, COX6, and SDHB), together with increased abundance of oxidative lipid species in severe-EMS-sEVs **(Fig. S7a-i; Fig. 4h, i, q, r**), we next investigated whether EMS-sEVs specifically localize to the mitochondria following uptake by HUtMECs. Using the same live-cell imaging workflow described above, cells were treated with MemGlow™-labeled mild- or severe-EMS-sEVs and monitored over a 10 h time course. To capture early intracellular uptake dynamics, additional early timepoints were included for assessment of sEV localization relative to mitochondria (**Fig. 7g, h**). To specifically assess potential mitochondrial association of internalized sEVs, BODIPY TR staining was replaced with a mitochondria-targeted fluorescent stain. Both treatment groups demonstrated progressive co-localization of MemGlow™ fluorescence with MitoTracker™ Deep Red staining over time (**Fig. 7e, f**), enabling assessment of sEV-associated fluorescence relative to mitochondrial structures. Quantitative co-localization analysis demonstrated significantly greater mitochondrial-associated fluorescence in severe EMS-sEV-treated cells compared with mild-EMS-sEV-treated cells between 4 and 10 h post-treatment (T2–T5; **Fig. 7g, h**). Together, these findings demonstrate efficient uptake and mitochondrial localization of EMS-sEVs by HUtMECs, with enhanced mitochondrial association of severe-EMS-sEVs, reflecting increased capacity to modulate endothelial cell function.

To determine whether uptake of lesion-derived sEVs translated into functional alterations in endothelial cell behaviour, proliferation, apoptosis, and cytokine secretion were assessed following treatment with the same cohort of stage- and tissue-specific sEV populations. Mild-EMS-sEVs induced a significant increase in HUtMEC proliferation following 12 h incubation relative to control conditions (**Fig. 8a**), consistent with enhanced endothelial metabolic activity and proliferative potential. In contrast, no significant differences in apoptosis (measured via Caspase 3/7 activity) were observed across treatment groups during the same timeline (**Fig. 8b**), suggesting that sEV exposure did not substantially alter apoptotic signaling under these experimental conditions.

**Figure 8.**
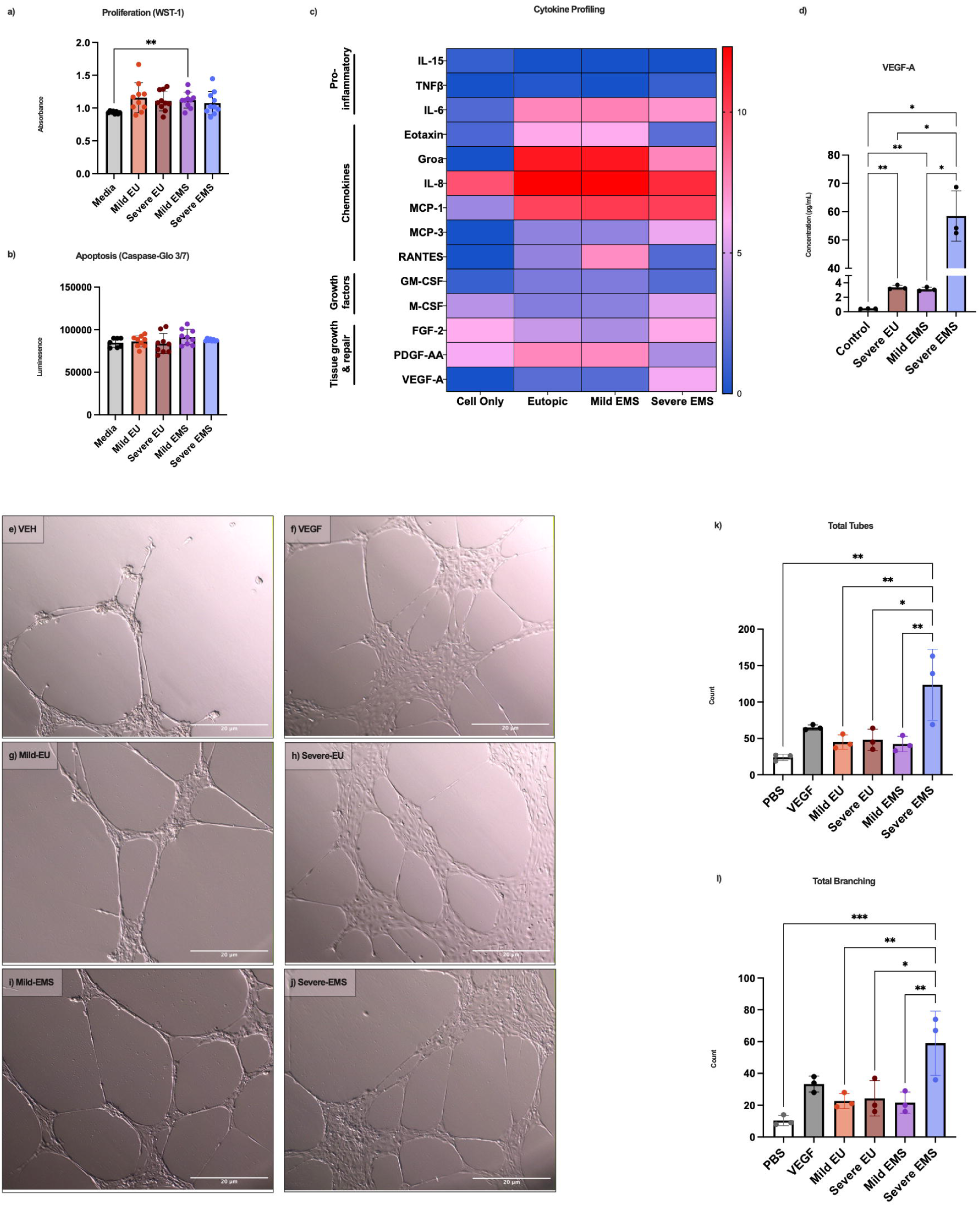
EM tissue-derived sEVs induce stage- and tissue-specific functional alterations in HUtMECs. HUtMECs were treated with pooled sEVs (n=3) isolated from matched EU and EMS tissues of mild- and severe-EM patients. Functional assays were performed following 12 h incubation unless otherwise indicated. **(A)** Treatment of HUtMECs with sEVs derived from mild EMS significantly increased cellular proliferation relative to severe EMS-derived sEVs and matched EU-derived sEVs, as determined by WST-1 assay. (**B)** No significant differences in apoptosis were observed between experimental groups, as demonstrated by comparable Caspase-Gio 3/7 activity. **(C)** Log-transformed heatmap analysis of conditioned media from HUtMECs treated with pooled EU- and EMS-derived sEVs from mild and severe EM patients demonstrated increased expression of chemokines, inflammatory mediators, and growth and repair factors in cells treated with EMS-derived sEVs relative to EU and untreated control groups. **(D)** HUtMECs treated with severe EMS-derived sEVs expressed significantly increased levels of VEGF-A relative to vehicle (PBS), EU-sEV and mild EMS-sEV treated cells following 12 h incubation. **(E-M)** Treatment of HUtMECs with sEVs derived from mild and severe EMS and matched EU induced endothelial tubulogenesis following 12 h incubation. **(E-J)** HUtMECs demonstrated stage- and tissue-specific differences in tube formation. Representative images were acquired using the Leica MICA at 10x magnification. Scale bar=20 pm. (**K-L)** Quantitative analysis of tube formation revealed that HUtMECs treated with severe EMS-derived sEVs exhibited significantly increased **(K)** total tube number and **(L)** total branching nodes relative to mild EU-derived sEVs, mild EMS-derived sEVs, severe EU-derived sEVs, and PBS-treated controls. Tube formation analysis was performed using WimTube software (WIMASIS). Statistical analysis was conducted using one-way ANOVA, withTukey’s multiple comparisons test (*P < 0.05, **P < 0.01, ****P< 0.0001).

Analysis of conditioned media further revealed distinct cytokine release profiles following treatment with stage- and tissue-specific sEV populations. Severe-EMS-sEVs induced significantly greater secretion of chemokines, and growth- and tissue repair-associated factors compared with other treatment groups (**Fig. 8c**). MCP-1 secretion was significantly increased relative to cell-only controls (P < 0.05), whereas MCP-3 (P < 0.05), FGF-2 and VEGF-A (P < 0.05), and M-CSF (P < 0.01) were significantly elevated compared with cell-only controls, EU-sEVs, and mild-EMS-sEVs. (**Fig. 8c**). In contrast, mild-EMS- and severe-EU-sEVs induced significantly greater secretion of pro-inflammatory cytokines and chemokines (IL-6, IL-8; P < 0.05, GROα; P < 0.01), with a concomitant increase in PDGF-AA (P < 0.01) relative to other treatment groups, suggesting potential involvement in endothelial and environmental remodeling programs linked to lesion establishment and vascular adaptation (**Fig. 8c**).

Building on the increased VEGF-A secretion observed following severe-EMS-sEV treatment (**Fig. 8d**), an angiogenic tube formation assay was performed to determine whether sEV-induced endothelial signaling translated into functional changes in angiogenic capacity. Treatment with mild-EMS- and severe-EMS-, as well as matched EU-sEVs, induced endothelial tubulogenesis following 12 h incubation (**Fig. 8e-j**). Live-cell imaging revealed more extensive vascular network formation in severe-EMS sEV-treated HUtMECs (**Fig. 8j**) relative to vehicle control, mild sEV- and severe-EU sEV-treated HUtMECs(**Fig. 8e, g-i)**, with a morphology that appeared comparable to VEGF-A-treated positive control conditions (**Fig. 8f**). These qualitative observations were supported by quantitative WimTube analysis demonstrating significant increases in total tube number and branching points (**Fig. 8k-m**). These results demonstrate that sEVs derived from distinct disease stages and tissue sources differentially modulate endothelial cell uptake, cytokine secretion, and angiogenic behaviour.

## 4. Discussion

EM remains a heterogeneous disease with substantial impacts on quality of life, reproductive health, and clinical management in over ∼200M women worldwide.^24^ Current therapeutic strategies, including hormonal suppression and surgical intervention, primarily focus on symptom management and lesion removal, but do not prevent disease recurrence and require careful consideration in individuals seeking to preserve or achieve fertility.^5,25^ Despite its considerable clinical burden, the molecular mechanisms governing lesion establishment, ectopic tissue survival, and progression toward chronic inflammatory, vascular, and fibrotic states remain incompletely understood. In particular, the early molecular events that enable refluxed endometrial tissue to implant and persist within the peritoneal cavity remain poorly defined.^1,2^ This knowledge gap is further complicated by the substantial biological heterogeneity of EM, where distinct lesion subtypes exhibit diverse molecular and cellular features despite frequent classification by rASRM stage. Together, these challenges contribute to delayed diagnosis, limited non-invasive biomarkers, and a lack of mechanism-directed therapeutic strategies. Previous studies have shown that sEVs are altered in EM and contribute to intercellular communication through the transfer of proteins, lipids, and nucleic acids that reflect the physiological state of their cells of origin. ^6,7^ These findings have highlighted sEVs as both mediators of disease biology and promising sources of non-invasive biomarkers. However, most studies have examined individual cargo classes or isolated biological compartments, limiting our understanding of stage- and tissue-specific sEV signatures.

In the present study, we performed a comprehensive stage- and tissue-specific characterization of EM-derived sEVs isolated from eutopic endometrium, ectopic lesions, PF, and plasma using complementary surface immune phenotyping, proteomic, lipidomic, and functional analyses. Together, these approaches identified coordinated molecular programs associated with immune remodeling, epithelial plasticity, ECM organization, metabolic adaptation, and vascular signaling that evolve throughout disease progression. Importantly, functional studies demonstrated that disease-associated sEV populations actively influence endothelial cell behaviour *in vitro*, supporting a role for sEV-mediated intercellular communication in shaping the endometriotic lesion microenvironment. Collectively, these findings provide new insight into the molecular mechanisms underlying disease heterogeneity and establish an integrated framework for understanding how stage- and tissue-specific alterations in sEV composition may contribute to EM pathogenesis.

Characterization of sEV populations confirmed expected morphology and size distributions across all sample types, while revealing significant stage- and source-specific differences in vesicle properties. TEM analysis revealed increased structural heterogeneity within severe-PF- and EMS-derived sEV preparations, including distinct intravesicular electron-dense structures. While the composition of these structures remains unknown, their increased prevalence in severe disease-associated sEV populations suggests that disease stage is accompanied by alterations in vesicle organization beyond changes in size or abundance. Severe-stage plasma contained increased concentrations of circulating sEVs together with selective enrichment of CD9 expression, consistent with previous reports identifying CD9 as a predominant marker of circulating biofluid-derived EV populations.^26^ In contrast, both severe-stage EMS and EU tissue-derived sEVs demonstrated increased expression of CD63 and CD81, supporting stage-dependent remodeling of local vesicle populations and aligning with previous studies reporting preferential enrichment of these tetraspanins in tissue-derived EV samples.^27^ Together, these findings highlight that canonical EV markers are highly context-dependent and emphasize the importance of considering sample origin when interpreting sEV-associated signatures in EM.

Surface phenotyping further demonstrated that sEV populations undergo pronounced stage- and tissue-dependent phenotypic remodeling, indicating that severe disease is accompanied not only by quantitative changes in vesicle abundance but also by alterations in molecular composition that may influence recipient cell interactions. While circulating plasma-derived sEVs retained relatively conserved surface profiles, lesion-derived vesicles exhibited disease stage-associated enrichment of immune-, adhesion-, and antigen presentation-associated markers, suggesting increasing specialization of local sEV populations within the inflammatory lesion microenvironment. Importantly, these surface signatures may be interpreted as indicators of altered sEV composition within the lesion microenvironment, which could arise from changes in cargo sorting, cellular composition, or the activation state of EV-producing cells.

Severe-stage lesions were accompanied by reduced epithelial- and stemness-associated markers, including CD133/1 and EpCAM (CD326), together with increased HLA-II expression. Given the established roles of EpCAM in epithelial organization, these findings suggest a shift in lesion-derived sEV surface composition away from epithelial-associated features toward immune-associated signatures in severe EM. However, these changes may reflect both altered molecular sorting into sEVs and broader remodeling of the lesion cellular landscape, where increased stromal activation, ECM remodeling, and immune infiltration may alter the relative contribution of epithelial-, stromal-, and immune-derived vesicle populations.^28–30^ Although CD133 expression varies depending on biological source and analytical approach, CD133/prominin-1 has been associated with epithelial and stemness features in endometrial and endometriotic tissues, and recent organoid-derived EM studies demonstrate incorporation of CD133 into epithelial-derived sEV populations.^18,27^ Our findings extend these observations by demonstrating stage-dependent reductions in sEV-associated CD133/1 across both lesion- and PF-derived sEVs, suggesting that alterations in epithelial-associated sEV signatures within the local lesion microenvironment are associated with severe disease. Although systemic surface phenotypes were comparatively subtle, severe-plasma-derived sEVs demonstrated increased expression of adhesion- and platelet-associated markers (CD29, CD41b, CD42a, and CD62P), consistent with growing evidence linking platelet activation, vascular remodeling, and coagulation pathways to EM pathophysiology.^28,31,32^ Together with the observed enrichment of CD9 in severe-plasma-sEVs, these alterations highlight the context-dependent nature of circulating sEV surface signatures and support the utility of plasma-derived vesicles as minimally invasive indicators of EM-associated remodeling, providing a potential framework for future biomarker development and patient stratification.

Importantly, these stage- and tissue-specific surface phenotypes closely paralleled the molecular programs identified through proteomic and lipidomic profiling, revealing coordinated yet compartment-specific remodeling of sEV composition across EM. Tissue-derived sEV proteomes demonstrated distinct stage-associated molecular programs, with mild-stage sEVs enriched for signatures associated with epithelial maintenance, adhesion stability, and proliferative remodeling, while severe-stage sEVs exhibited increased representation of immune activation, ECM remodeling, vascular signaling, and inflammatory stress adaptation. The increased structural heterogeneity observed by TEM in severe disease-associated sEV populations further supports the concept that severe EM is accompanied by coordinated remodeling of sEV composition and organization. These structural differences coincided with distinct lipidomic signatures in EMS-derived sEVs, suggesting that alterations in membrane composition may influence vesicle organization, cargo distribution, or biophysical properties. In contrast to EMS-derived sEVs, plasma-sEVs exhibited a largely shared disease-associated lipid signature across mild- and severe-stage EM relative to healthy controls, with additional stage-associated lipid alterations observed in severe disease. Notably, LPC(16:0), oxidized triglycerides, and glycosphingolipids were recurrently altered across multiple biological compartments, suggesting that coordinated membrane remodeling and oxidative lipid metabolism represent conserved features of EM-derived sEVs despite broader tissue- and stage-specific differences. Together, alterations in surface markers, structural features, and molecular cargo demonstrate that disease-associated sEV remodeling extends beyond individual markers or cargo classes, reflecting broader restructuring of vesicle biology across disease compartments.

To our knowledge, this represents the first comprehensive integration of sEV-associated proteomic and lipidomic profiles across distinct sample types and disease stages in EM. Integrated multi-omic analyses revealed coordinated relationships between complementary molecular cargo classes, demonstrating that stage- and tissue-dependent alterations identified through individual datasets converge on shared patterns of sEV remodeling. These findings suggest that local lesion-associated vesicles undergo more tightly coupled molecular remodeling, while circulating sEV populations retain broader disease-associated signatures shaped by diverse cellular contributions. Notably, these multi-omic signatures further align with previously identified regulatory RNA alterations in EM-derived sEVs, suggesting that disease-associated remodeling extends across multiple cargo classes rather than representing isolated changes within individual molecular layers. Lesion-derived vesicles exhibited altered expression of let-7 family members, miR-23a, miR-206, and miR-320a, together with dysregulation of the long non-coding RNAs H19 and NEAT1, while plasma-derived sEVs displayed distinct circulating miRNA profiles relative to healthy controls.^13^ These regulatory RNA programs have been implicated in biological processes central to EM progression, including inflammatory signaling, cellular plasticity, invasive phenotypes, and tissue remodeling.^33–36^ The convergence of RNA, protein, and lipid signatures supports a model in which distinct sEV cargo classes collectively contribute to disease-associated extracellular signaling and highlights the value of integrated molecular profiling for defining conserved and tissue-specific mechanisms underlying EM pathogenesis.

To determine whether these coordinated molecular programs translated into biologically meaningful effects on recipient cells, we evaluated the functional effects of tissue-derived sEVs on HUtMECs, an endothelial cell model representative of vascular remodeling during lesion establishment and persistence. Given the dependence of ectopic lesions on vascularization, endothelial cells represent a key cellular target through which EM-derived sEVs may influence lesion development. Although both mild- and severe-EMS-sEVs were efficiently internalized, severe-stage vesicles exhibited significantly greater intracellular accumulation and preferential mitochondrial localization, indicating that disease stage influences not only sEV molecular cargo but also the dynamics of recipient cell interactions. This observation is of particular interest given the enrichment of mitochondrial-associated proteins (ATP5F1A-C, NDUF6,8,10, COX4I1, COX6, and SDHB) and oxidative lipid species (PEt37:2+4O, Cer d44:1+O, Cer t42:0+O) within severe-EMS-derived sEVs.

Additionally, the reduced abundance of CoQ10 in plasma-derived sEVs from individuals with severe EM suggests that mitochondrial- and oxidative stress-associated alterations may extend beyond the lesion microenvironment into the systemic circulation. Metabolic dysregulation and oxidative stress are increasingly recognized features of EM pathophysiology, with previous metabolomic studies identifying alterations in energy metabolism and systemic metabolic profiles, alongside disruption of lipid metabolic pathways, including phospholipid-associated remodeling, in individuals with EM.^37,38^ Together, these findings suggest that sEV-associated lipid and protein remodeling reflects broader metabolic and oxidative adaptations associated with disease stage occurring both within the lesion microenvironment and systemically through circulating sEV populations. Moreover, the enhanced mitochondrial localization of severe-EMS-derived sEVs in recipient endothelial cells raises the possibility that EM-derived sEVs may actively influence mitochondrial-associated processes through the transfer of bioactive cargo. This concept is supported by growing evidence that extracellular vesicles selectively package and transfer mitochondrial-associated proteins capable of influencing recipient cell metabolism and bioenergetic homeostasis.^39^ Future studies will be required to determine whether EM-derived sEVs directly alter mitochondrial function and metabolic activity within recipient cells.

Although circulating plasma-sEVs provide valuable insight into systemic disease-associated alterations, lesion-derived sEVs were selected for functional assessment because they directly reflect the local cellular environment in which ectopic lesions establish and undergo vascular remodeling. Importantly, the distinct endothelial responses induced by EMS-derived sEVs were not explained by differences in vesicle internalization across disease stages, highlighting the importance of stage-specific sEV cargo composition in shaping endothelial functional responses. The stage-specific effects observed in HUtMECs were consistent with the distinct molecular signatures identified within mild- and severe-stage sEV populations. Mild-EMS-derived sEVs preferentially promoted endothelial proliferation and induced cytokine programs associated with inflammatory activation (IL-6, IL-8, and GROa) and vascular adaptation (PDGF-AA) in mild EM, aligning with the enrichment of proteins associated with epithelial maintenance, remodeling, and proliferative signaling identified in mild-stage sEVs. In contrast, severe-EMS-derived sEVs promoted a more pronounced pro-angiogenic phenotype characterized by increased VEGF-A secretion, enhanced tube formation, and elevated expression of chemokines involved in immune recruitment and tissue remodeling (MCP-1, MCP-3, M-CSF, and FGF-2), consistent with the enrichment of inflammatory, vascular, and ECM-associated molecular programs identified in severe EM. Together, these findings suggest that stage-dependent sEV cargo composition contributes to distinct endothelial responses, highlighting the functional relevance of sEV molecular remodeling in EM pathogenesis.

Collectively, the functional phenotypes induced by lesion-derived sEVs closely mirrored the coordinated molecular programs identified through integrated surface phenotyping, proteomic, and lipidomic analyses, demonstrating that these molecular signatures reflect biologically relevant disease-associated processes rather than descriptive differences alone. These findings support a model in which ectopic lesion establishment and persistence are shaped by coordinated communication within a lesion-supportive microenvironment,^1–4^ with stage- and tissue-specific sEV populations coordinating immune activation, epithelial plasticity, vascular adaptation, ECM remodeling, and metabolic reprogramming in EM. By integrating complementary molecular layers with functional validation, this study provides a novel systems-level framework for understanding EM biology that would not be captured through individual omic approaches alone. Importantly, this work identifies lesion-derived sEVs as a previously underappreciated component of the EM microenvironment and highlights their potential as mechanistic mediators and sources of clinically relevant molecular signatures Although translation of patient-derived EM findings remains challenging due to substantial clinical and biological heterogeneity, including variation in lesion subtype, anatomical location, hormonal status, symptom severity, and frequence of comorbid conditions, larger clinically stratified cohorts will be essential to validate these signatures and define their utility across the diverse spectrum of EM. Ultimately, these findings provide a foundation for future studies aimed at leveraging integrated sEV-based molecular profiling to improve disease classification, uncover clinically relevant biomarkers, and advance precision approaches for EM management.

## Author Contributions

J.P.H conceived and conducted experiments, analyzed data, and wrote the manuscript. K.B.Z. and D.J.S. assisted with experiments and processing human patient samples. D.H. isolated sEV samples for lipidomic analyses. O.B. and B.A.L. contributed human patient samples. C.T. conceived experiments, provided reagents and financial support. All authors read, edited, and approved the manuscript.

## Acknowledgements

We thank Oliver Jones for assistance with TEM sample processing and imaging, and Jeffrey Mewburn for valuable microscopy expertise and guidance. We also thank the Abraham laboratory for training and support with NTA and for access to their instrumentation. We thank Kira King and Jessica Pudwell for their assistance with patient sample coordination. We thank Creative Proteomics and the Proteomics and Molecular Analysis Core at the Research Institute of the McGill University Health Centre (RI-MUHC) for their sequencing services.

## REFERENCES

1. Zondervan, K. T., Becker, C. M. & Missmer, S. A. Endometriosis. New England Journal of Medicine 382, 1244–1256 (2020).

2. As-Sanie, S. et al. Endometriosis. JAMA 334, 64 (2025).

3. Horne, A. W. & Missmer, S. A. Pathophysiology, diagnosis, and management of endometriosis. BMJ 379, e070750 (2022).

4. Symons, L. K. et al. The Immunopathophysiology of Endometriosis. Trends Mol. Med. 24, 748–762 (2018).

5. Agarwal, S. K. et al. Clinical diagnosis of endometriosis: a call to action. Am. J. Obstet. Gynecol. 220, 354.e1–354.e12 (2019).

6. van Niel, G. et al. Challenges and directions in studying cell–cell communication by extracellular vesicles. Nat. Rev. Mol. Cell Biol. 23, 369–382 (2022).

7. Yáñez-Mó, M., et al. Biological properties of extracellular vesicles and their physiological functions. J. Extracell. Vesicles 4, (2015).

8. Beal, J. R., Ma, Q., Bagchi, I. C. & Bagchi, M. K. Role of Endometrial Extracellular Vesicles in Mediating Cell-to-Cell Communication in the Uterus: A Review. Cells 12, 2584 (2023).

9. Matsuura, Y. et al. Exosomal miR-155 Derived from Hepatocellular Carcinoma Cells Under Hypoxia Promotes Angiogenesis in Endothelial Cells. Dig. Dis. Sci. 64, 792–802 (2019).

10. Bertolini, I. et al. Small Extracellular Vesicle Regulation of Mitochondrial Dynamics Reprograms a Hypoxic Tumor Microenvironment. Dev. Cell 55, 163–177.e6 (2020).

11. Zhao, G. et al. Exosomal Sonic Hedgehog derived from cancer-associated fibroblasts promotes proliferation and migration of esophageal squamous cell carcinoma. Cancer Med. 9, 2500–2513 (2020).

12. Chu, X. et al. Extracellular vesicles in endometriosis: role and potential. Front. Endocrinol. (Lausanne). 15, (2024).

13. Khalaj, K., et al. Extracellular vesicles from endometriosis patients are characterized by a unique miRNA-lncRNA signature. JCI Insight 4, (2019).

14. Dai, L., Gu, L. & Di, W. MiR-199a attenuates endometrial stromal cell invasiveness through suppression of the IKK /NF-B pathway and reduced interleukin-8 expression. Mol. Hum. Reprod. 18, 136–145 (2012).

15. Hawkins, S. M. et al. Functional MicroRNA Involved in Endometriosis. Molecular Endocrinology 25, 821–832 (2011).

16. Haikalis, M. E., Wessels, J. M., Leyland, N. A., Agarwal, S. K. & Foster, W. G. MicroRNA expression pattern differs depending on endometriosis lesion type†. Biol. Reprod. 98, 623–633 (2018).

17. Saare, M. et al. Challenges in endometriosis miRNA studies — From tissue heterogeneity to disease specific miRNAs. Biochimica et Biophysica Acta (BBA) - Molecular Basis of Disease 1863, 2282–2292 (2017).

18. Wang, Y., et al. Human Endometriotic Lesion-Derived Small Extracellular Vesicles Impair Macrophage Function in the Peritoneal Microenvironment. J. Extracell. Vesicles 15, (2026).

19. Sisnett, D. J. et al. An investigation of the IL-23/Th17 axis and transcriptomic profiles of T helper subsets in endometriosis. Preprint at 10.64898/2026.06.11.731688 (2026).

20. Saint-Pol, J. & Culot, M. Minimum information for studies of extracellular vesicles (MISEV) as toolbox for rigorous, reproducible and homogeneous studies on extracellular vesicles. Toxicology in Vitro 106, 106049 (2025).

21. Monash Proteomics and Metabolomics Platform & Monash Bioinformatics Platform, Monash University. DIA-Analyst: An interactive web-platform to analyze and visualize proteomics data preprocessed with Spectronaut and DIA-NN. https://analyst-suites.org/apps/dia-analyst/.

22. Zhou, G., Ewald, J. & Xia, J. OmicsAnalyst: a comprehensive web-based platform for visual analytics of multi-omics data. Nucleic Acids Res. 49, W476–W482 (2021).

23. Ewald, J. D. et al. Web-based multi-omics integration using the Analyst software suite. Nat. Protoc. 19, 1467–1497 (2024).

24. World Health Organization (WHO). Endometriosis. Fact sheets. https://www.who.int/news-room/fact-sheets/detail/endometriosis. (2023).

25. Giudice, L. C. & Kao, L. C. Endometriosis. The Lancet 364, 1789–1799 (2004).

26. Karimi, N., Dalirfardouei, R., Dias, T., Lötvall, J. & Lässer, C. Tetraspanins distinguish separate extracellular vesicle subpopulations in human serum and plasma – Contributions of platelet extracellular vesicles in plasma samples. J. Extracell. Vesicles 11, (2022).

27. Garcia-Martin, R., Brandao, B. B., Thomou, T., Altindis, E. & Kahn, C. R. Tissue differences in the exosomal/small extracellular vesicle proteome and their potential as indicators of altered tissue metabolism. Cell Rep. 38, 110277 (2022).

28. Zhang, Q., Duan, J., Liu, X. & Guo, S.-W. Platelets drive smooth muscle metaplasia and fibrogenesis in endometriosis through epithelial–mesenchymal transition and fibroblast-to-myofibroblast transdifferentiation. Mol. Cell. Endocrinol. 428, 1–16 (2016).

29. Vissers, G., Giacomozzi, M., Verdurmen, W., Peek, R. & Nap, A. The role of fibrosis in endometriosis: a systematic review. Hum. Reprod. Update 30, 706–750 (2024).

30. Symons, L. K. et al. The Immunopathophysiology of Endometriosis. Trends Mol. Med. 24, 748–762 (2018).

31. Bortot, B. et al. Platelets as key cells in endometriosis patients: Insights from small extracellular vesicles in peritoneal fluid and endometriotic lesions analysis. The FASEB Journal 38, (2024).

32. Ding, D., Liu, X., Duan, J. & Guo, S.-W. Platelets are an unindicted culprit in the development of endometriosis: clinical and experimental evidence. Human Reproduction 30, 812–832 (2015).

33. Farzaneh, M. et al. Emerging roles of the long non-coding RNA NEAT1 in gynecologic cancers. J. Cell Commun. Signal. 17, 531–547 (2023).

34. Yuan, D. et al. Expression of lncRNA NEAT1 in endometriosis and its biological functions in ectopic endometrial cells as mediated via miR-124-3p. Genes Genomics 44, 527–537 (2022).

35. Ghazal, S. et al. H19 lncRNA alters stromal cell growth via IGF signaling in the endometrium of women with endometriosis. EMBO Mol. Med. 7, 996–1003 (2015).

36. Raveh, E., Matouk, I. J., Gilon, M. & Hochberg, A. The H19 Long non-coding RNA in cancer initiation, progression and metastasis – a proposed unifying theory. Mol. Cancer 14, 184 (2015).

37. Dai, Y. et al. Integrative analysis of transcriptomic and metabolomic profiles reveals abnormal phosphatidylinositol metabolism in follicles from endometriosis-associated infertility patients. J. Pathol. 260, 248–260 (2023).

38. Dutta, M. et al. Metabolomics reveals perturbations in endometrium and serum of minimal and mild endometriosis. Sci. Rep. 8, 6466 (2018).

39. Todkar, K. et al. Selective packaging of mitochondrial proteins into extracellular vesicles prevents the release of mitochondrial DAMPs. Nat. Commun. 12, 1971 (2021).

