## Supplemental Figures for "Endometriosis patient-derived small extracellular vesicles carry unique immune, proteomic and lipidomic signatures associated with mild and severe endometriosis"

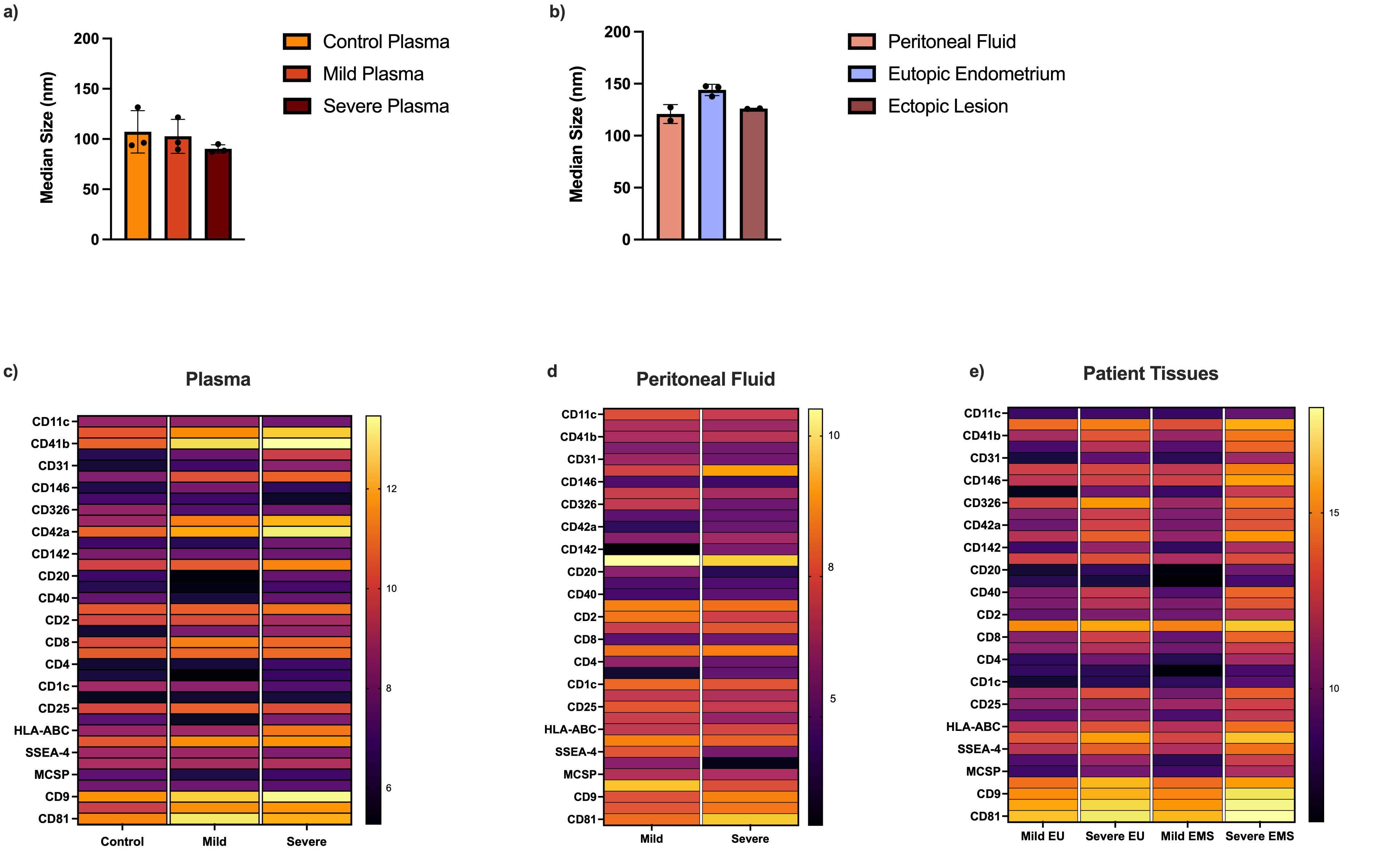


**Figure S1. Characterization and surface marker profiling of endometriosis (EM)-derived sEVs reveal stage- and tissue-specific differences in immune- and adhesion-related phenotypes. (A)** Median particle size distribution of plasma-derived sEVs from patients with mild EM, severe EM, and healthy controls (n = 3/ group). **(B)** Median particle size distribution of matched ectopic lesion (EMS)-, eutopic endometrium (EU)-, and peritoneal fluid (PF)-derived sEVs from EM patients (n = 3/ group). **(C-E)** Heatmap analysis of relative surface marker abundance normalized to bead-only controls demonstrated consistently high tetraspanin expression and low isotype control signal across all sample types. Statistical analyses were performed using one-way ANOVA with Tukey’s multiple comparisons test.

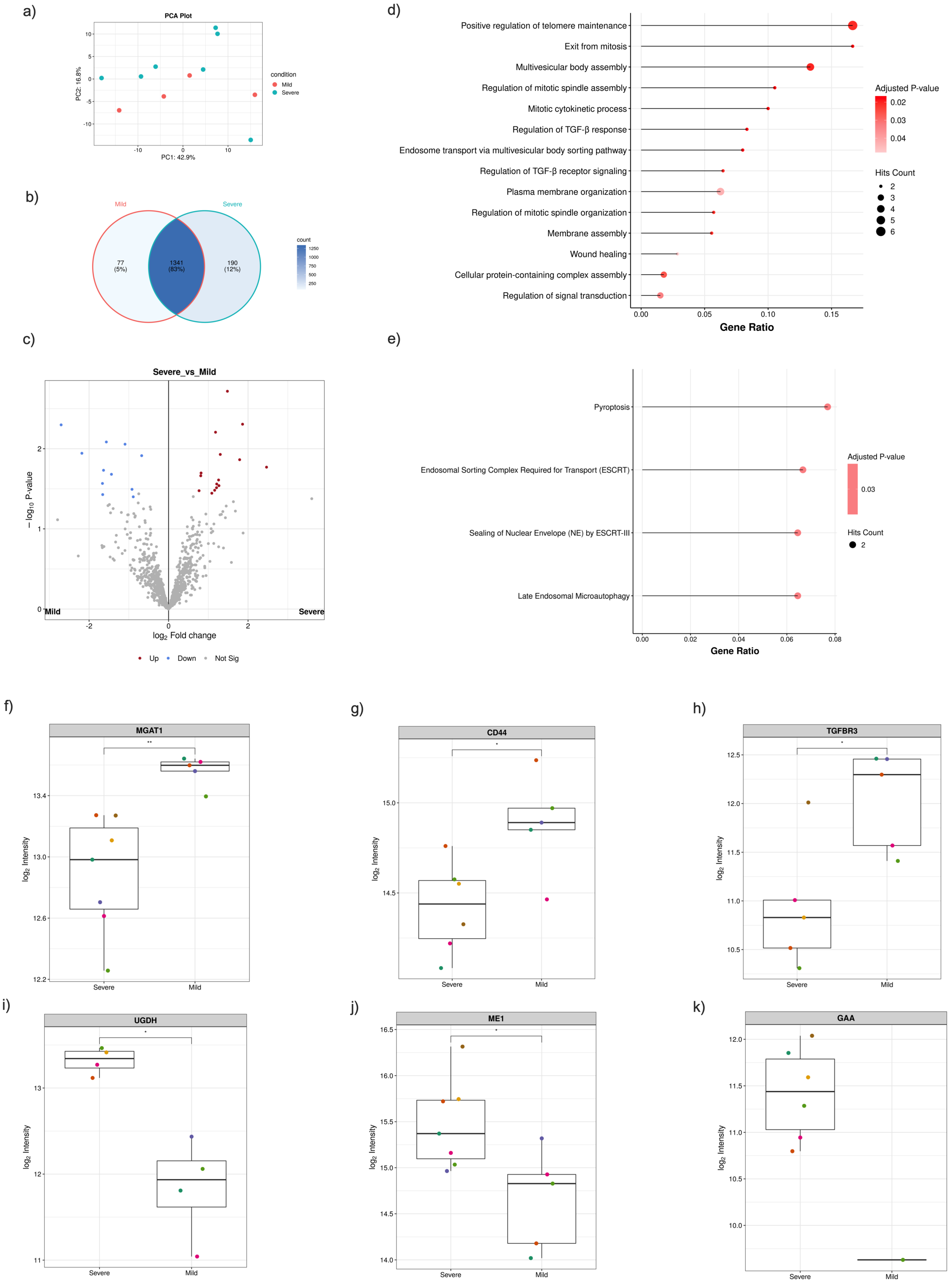


**Figure S2. Proteomic profiling of PF-derived sEVs reveals stage-associated alterations in immune signaling, vesicle trafficking, and metabolic remodeling in EM. (A)** Principal component analysis (PCA) of PF-derived sEV proteomes from mild- and severe-stage EM patients demonstrated partial clustering by disease stage. **(B)** Venn diagram analysis identified both shared and stage-specific protein expression patterns **(C)** Volcano plot identified significantly differentially expressed proteins between mild- and severe-stage PF-derived sEVs. Differential expression analysis was performed using a Benjamini–Hochberg adjusted P-value cutoff of 0.05 and log2 fold-change threshold ≥ 0.5 following variance stabilizing normalization (VSN), without data imputation. **(D-E)** Functional enrichment analyses revealed distinct biological pathways associated with disease stage progression in PF-derived sEVs. **(D)** Gene Ontology (GO) Biological Process (BP) enrichment analysis of mild versus severe PF-derived sEVs **(E)** Reactome pathway analysis demonstrated enrichment of mild versus severe PF-derived sEVs. Functional enrichment analyses were performed using over-representation analysis (ORA) with an adjusted P-value cutoff of 0.05. (**F-K)** Box plot analyses highlight significantly dysregulated proteins associated with immune modulation, ECM interactions, metabolic adaptation, and lesion-supportive signaling within PF-derived sEVs. **(F)** MGAT1, **(G)** CD44, **(H)** TGFBR3, **(I)** UGDH, **(J)** ME1, and **(K)** GAA expression.

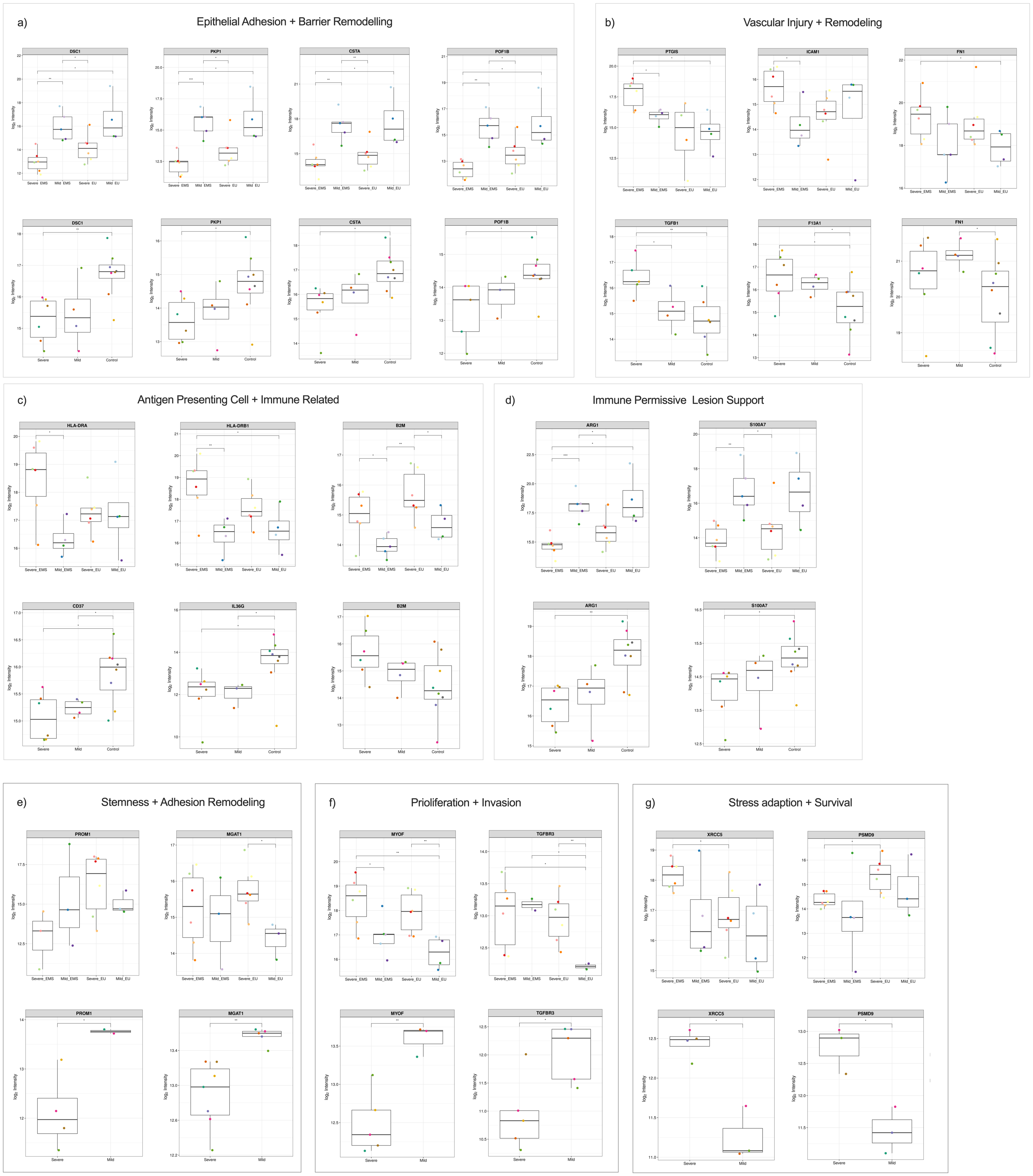


**Figure S3. Conserved axes between sEVs across EM-patient samples (A-D)** Relative abundance of proteins identified by proteomic profiling in mild- and severe-stage EU-, EMS-, and plasma-derived-sEVs box plots demonstrate increased abundance of **(A)** proteins associated with epithelial adhesion (DSC1, PKP1) and epithelial homeostasis (CSTA, POF1B) in healthy control plasma and mild-stage tissues, **(B)** inflammatory and remodeling-associated (TGFB1, FN1, ICAM1, and F13A1) across severe-stage plasma and lesion-derived sEVs, **(C)** antigen-presentation and inflammatory activation markers (HLA-related proteins, B2M) enriched in severe-stage tissues and plasma-derived sEVs, whereas immune homeostasis and neutrophil-associated proteins (CD37, IL36G) were relatively enriched in healthy controls, **(D** immune-permissive lesion support (ATG1 and S100A7) were enriched in mild-stage lesion-derived sEVs and healthy control plasma relative to severe-stage sEVs. **(E-G)** Relative abundance of proteins identified by proteomic profiling in mild- and severe-stage EU, EMS, and PF-derived-sEVs Box plots demonstrate increased abundance of **(E)** remodeling-associated proteins (CD133, MGAT1), **(F)** proliferative-associated processes (MYOF, TGFBR3) were preferentially enriched in mild-stage PF-derived sEVs, whereas reduced abundance was observed in corresponding tissue-derived populations, **(G)** proteins associated with inflammatory stress adaptation and proteostatic regulation (XRCC5, PSMD9) demonstrated concordant enrichment across severe-stage PF- and tissue-derived sEVs. Differential expression analysis was performed using a Benjamini–Hochberg adjusted p-value cutoff of 0.05 and log2 fold-change threshold ≥ 0.5 following variance stabilizing normalization (VSN), without data imputation.

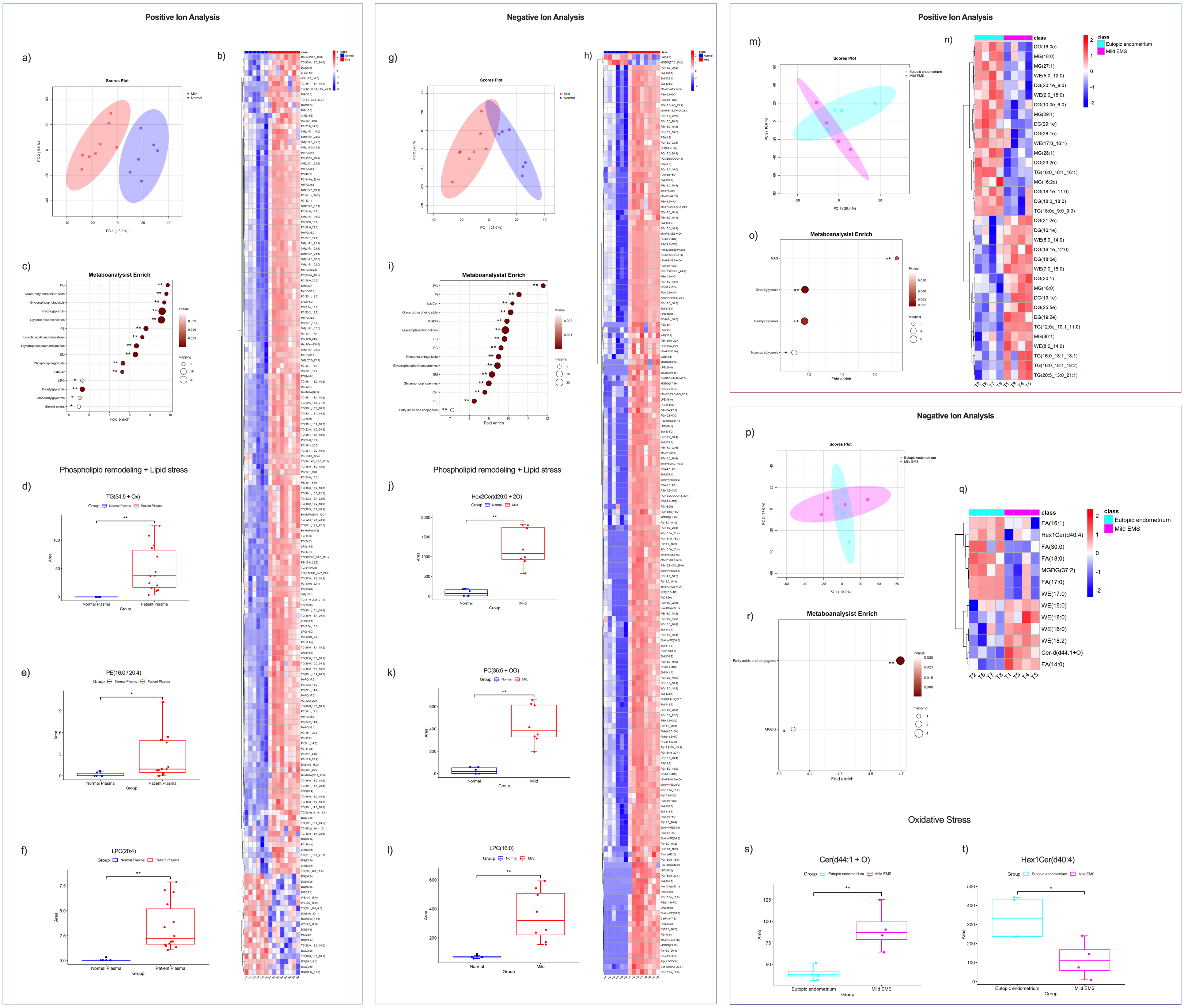


**Figure S4. Mild-stage derived sEV lipid profiles exhibit coordinated phospholipid remodeling and systemic oxidative stress–associated lipid signatures in EM. (A-F)** Positive ion analysis mode. **(A)** PCA demonstrated separation between mild-stage and control plasma sEV-derived lipid profiles, indicative of systemic lipid remodeling **(B)** Heatmap analysis revealed clear separation between mild-stage and control plasma samples, supporting lipidomic divergence associated with disease presence. **(C)** Dot plot enrichment analysis demonstrated over-representation of PG, TG, GPC, PE, SM, and CER in mild-plasma samples. **(D-F)** Representative box blots demonstrated that mild-plasma sEVs exhibited increased abundance of **(D)** TG(54:5 + O), **(E)** PE(16:0 / 20:4), and **(F)** LPC(16:0) relative to control plasma. **(G-I)** Negative ion analysis mode. **(G)** PCA demonstrated separation between mild-stage and control plasma sEV-derived lipid profiles **(H)** Heatmap analysis revealed clear separation between mild-stage and control plasma samples. **(I)** Dot plot enrichment analysis demonstrated over-representation of PG, PI, Cer, GPC, OS, and MGDG in mild-plasma samples. **(J-L)** Representative box blots demonstrated that mild-plasma exhibited increased abundance of **(J)** Hex2Cer(d29:0 + 2O), **(K)** PC(36:6 + OO), and **(L)** LPC(16:0), relative to control plasma. **(M-O)** Positive ion analysis mode. **(M)** PCA demonstrated partial separation between mild-EMS and EU sEV-derived lipid profiles. **(N)** Heatmap revealed separation between mild-EMS and EU samples, supporting lipidomic remodeling associated with tissue-compartment. **(O)** Dot plot enrichment analysis demonstrated over-representation of MAG, DG, and TG in mild-EMS samples. **(P-T)** Negative ion analysis mode. **(P)** PCA demonstrated partial separation between mild-EMS and EU sEV-derived lipid profiles. **(Q)** Heatmap analysis revealed separation between mild-EMS and EU samples. **(S)** Dot plot enrichment analysis demonstrated over-representation of FA and MGDG in mild-EMS samples. **(T-U)** Representative box blots demonstrated that mild-EMS exhibited increased abundance of **(S)** Cer(d44:1 + O) and **(T)** Hex1Cer(d40:4) relative to EU samples. Analysis was filtered and confirmed by combining the results of the VIP values (VIP >1.5), fold-change (|log2 FC| > 1) and t-test (*P < 0.05, **P < 0.01).

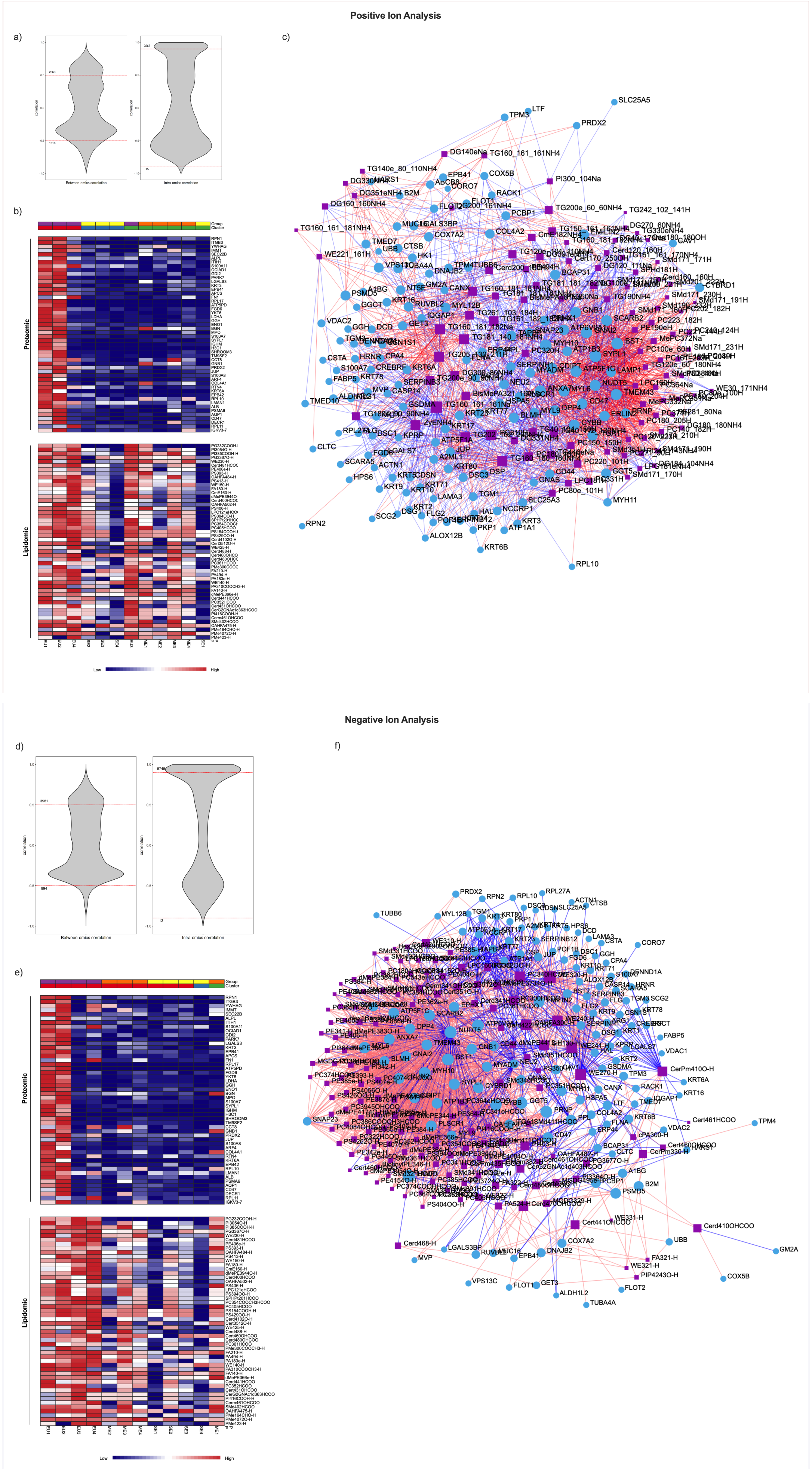


**Figure S5. Integrated proteomic and lipidomic correlation analysis identifies coordinated molecular relationships across tissue-derived sEVs. (A-C)** Positive ion tissue analysis. **(A)** Feature distribution plot of integrated proteomic and lipidomic datasets following preprocessing. **(B)** Unsupervised hierarchical clustering of integrated protein and lipid features demonstrated partial separation of mild- and severe-stage tissue-derived sEV samples, with limited overlap between groups. **(C)** Correlation network illustrating positive (red) and negative (blue) associations between protein (blue circles) and lipid (purple squares) features. **(D-F)** Negative ion tissue analysis. **(D)** Feature distribution plot. **(E)** Unsupervised hierarchical clustering demonstrated partial stage-associated separation of tissue-derived sEV samples. **(F)** Correlation network showing integrated protein-lipid relationships.

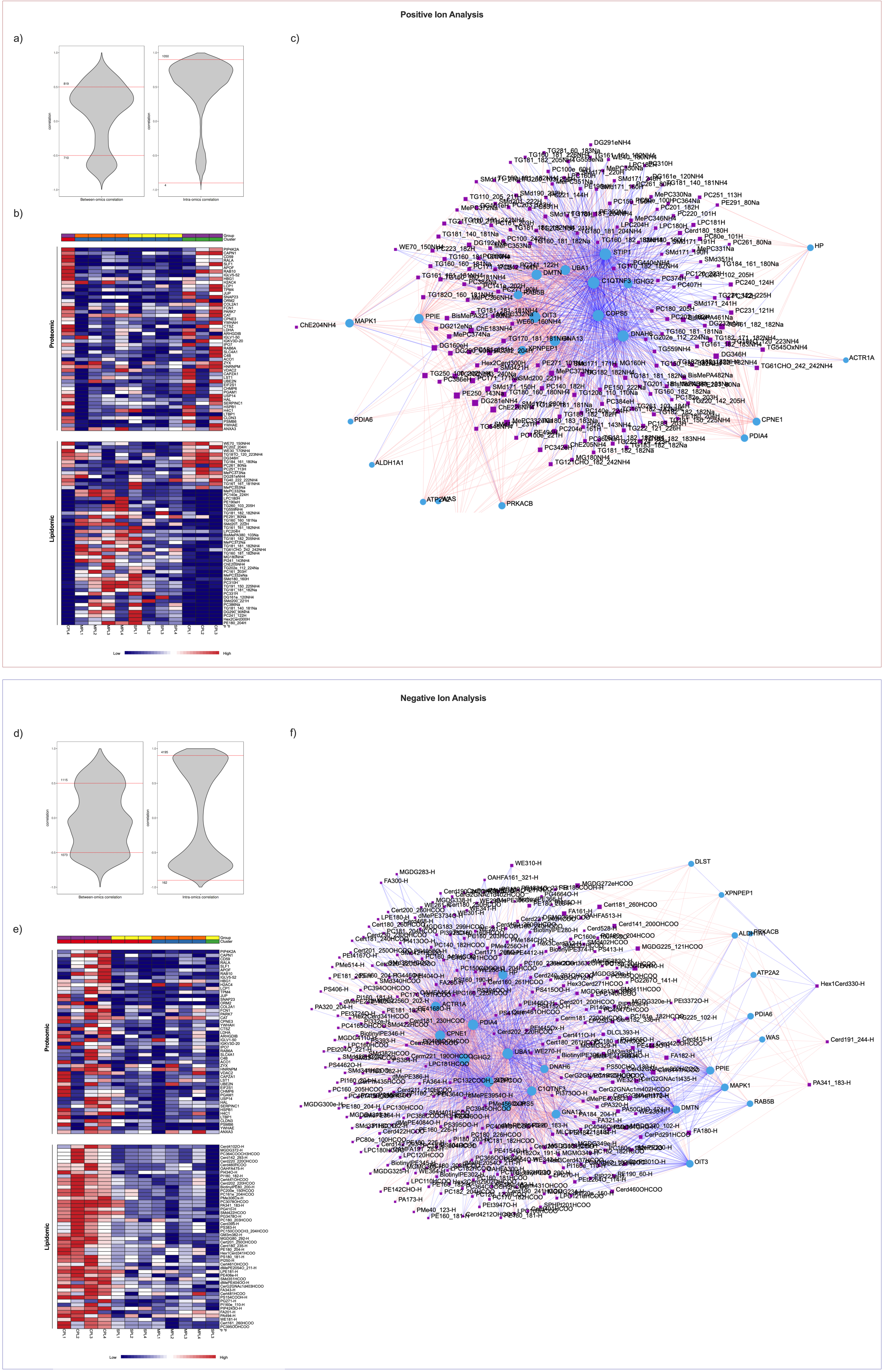


**Figure S6. Integrated proteomic and lipidomic correlation analysis identifies coordinated molecular relationships across plasma-derived sEVs. (A-C)** Positive ion plasma analysis. **(A)** Feature distribution plot. **(B)** Unsupervised hierarchical clustering demonstrated partial separation between control and EM plasma-derived sEV samples with limited overlap. **(C)** Correlation network showing positive and negative protein-lipid associations. **(D-F)** Negative ion plasma analysis. **(D)** Feature distribution plot. **(E)** Unsupervised hierarchical clustering demonstrated partial separation of plasma-derived sEV samples with modest overlap between disease groups. **(F)** Correlation network showing integrated protein-lipid relationships. For all network analyses, proteins are represented as blue circles, lipid species as purple squares, positive correlations as red edges, and negative correlations as blue edges.

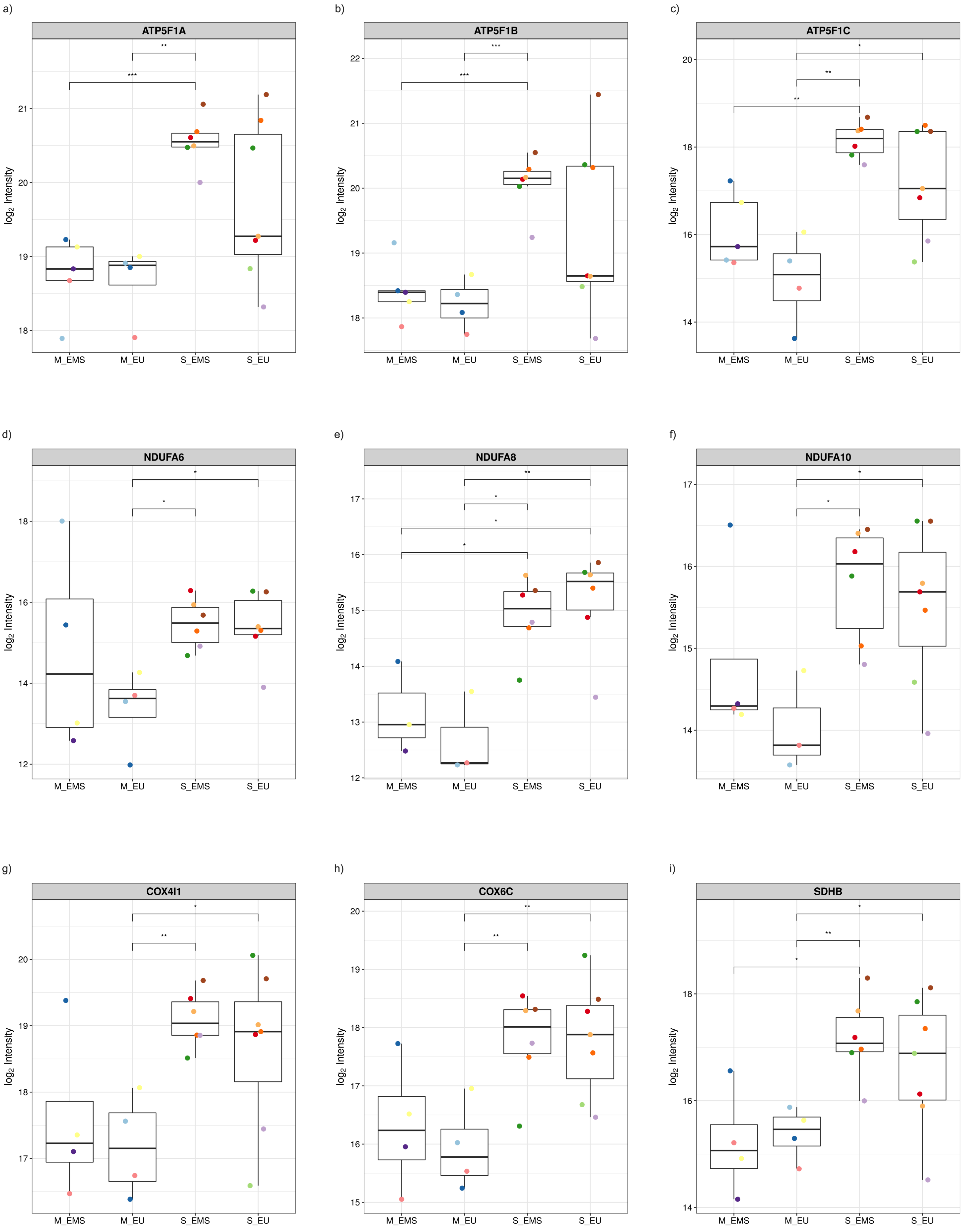


**Figure S7. Severe-EMS-derived sEVs exhibit enrichment of mitochondrial oxidative phosphorylation-associated proteins. (A-I)** Representative box plots demonstrated relative abundance of mitochondrial-associated proteins identified by proteomic profiling in mild- and severe-stage EU and EMS-derived sEVs. Box plots demonstrate increased abundance of **(A-C)** ATP synthase-associated proteins (ATP5F1A, ATP5F1B, and ATP5F1C), **(D-F)** electron transport chain components, including NADH dehydrogenase complex I subunits (NDUF6, NDUF8, NDUF10), **(G-H)** cytochrome c oxidase components (COX4I1 and COX6C), and **(I)** succinate dehydrogenase complex II subunit B (SDHB), particularly within severe-stage-derived sEVs. Differential expression analysis was performed using a Benjamini–Hochberg adjusted p-value cutoff of 0.05 and log2 fold-change threshold ≥ 0.5 following variance stabilizing normalization (VSN), without data imputation.
